# The protein interactome of the Neuron Specific Gene family (NSG1-3)

**DOI:** 10.64898/2025.12.07.692831

**Authors:** Antonio Serrano Rodriguez, Malene Overby, Upasna Srivastava, Praveen Chander, Liliana Vega, Sandy Wilson, Katherine E. Zychowski, Srikant Rangaraju, Heidi Kaastrup-Müeller, Jason P. Weick

## Abstract

The Neuron-Specific Gene (NSG) family members (NSG1-3) play critical and diverse roles in neuronal protein trafficking, but their precise molecular functions remain poorly understood. Here, we employed proximity-labeling proteomics to map the interactomes of each NSG protein. Unlabeled mass spectrometry identified over 1,000 significantly enriched interactors compared with a cytoplasmic control, revealing substantial overlap between NSG1 and NSG2, and a more divergent profile for NSG3. Gene ontology and KEGG pathway analysis confirmed established associations with glutamatergic synapses and endosomal trafficking, while also uncovering unexpected links to presynaptic machinery, inhibitory synapses, and endoplasmic reticulum-associated protein translation, particularly for NSG3. Reciprocal biotinylation patterns and co-immunoprecipitation revealed novel heteromeric complex formation between NSG1 and NSG2, with limited interactions involving NSG3. All three NSGs biotinylated core AMPA receptor subunits and auxiliary proteins, while NSG1 and NSG2 also associated with NMDA receptors, GABA receptor subunits, as well as multiple presynaptic proteins. Moreover, NSG1 and NSG2 specifically biotinylated components of multiunit tethering complexes including neuron-specific retromer, and biotinylated a preponderance of ADAM10 substrates, reinforcing their role in proteolytic processing. Finally, despite the relatively divergent interactomes of NSG1 and NSG2 compared to NSG3, all family members robustly biotinylated amyloid precursor protein (APP), suggesting possible synergistic or competitive interactions that could shape APP proteolytic processing and/or trafficking. Together, these data provide a comprehensive systems-level view of NSG protein interactions, establishing a molecular framework for future investigations into NSG-mediated neural plasticity and disease mechanisms.

## INTRODUCTION

Neurons are highly polarized cells with an array of unique subcellular compartments including axons, dendrites, presynaptic boutons and post-synaptic dendritic spines. This structural diversity demands precisely regulated systems for the production, trafficking, and localization of proteins across secretory, endocytic, and transcytotic pathways (Kennedy and Ehlers, 2006). Among the many proteins orchestrating these processes are members of the Neuron-Specific Gene (NSG) family—NSG1, NSG2, and NSG3. As their name indicates they are expressed almost exclusively in neurons, and localize to multiple subcellular organelles including the Golgi apparatus, early and late endosomes, and the plasma membrane (Muthusamy et al., 2015; Yap et al., 2017). Interestingly, NSG genes emerged relatively late in evolution, appearing specifically in vertebrates, a pattern that aligns with the emergence of complex neural architectures and higher-order cognitive abilities (Emes et al., 2008; Muthusamy et al., 2009). However, their highly distributed expression pattern along with their involvement in a host of trafficking and protein processing pathways makes their specific contributions to cellular function enigmatic.

Among their best-characterized functions is the regulation of glutamatergic signaling through control of AMPA receptor (AMPAR) trafficking. Disruption of NSG1 impairs AMPAR recycling following endocytosis and reduces long-term potentiation (LTP; Steiner et al., 2002; Alberi et al., 2005). Surprisingly, NSG1 knockout animals show no overt learning and memory deficits, instead exhibiting increased anxiety and motor coordination abnormalities (Austin et al., 2022). In contrast, NSG2 knockout mice demonstrate enhanced behavioral flexibility and more rapid acquisition in trace fear conditioning tasks (Zimmerman et al., 2024), despite the fact that NSG2 appears to promote surface AMPAR expression (Chander et al., 2019). NSG3 is essential for clathrin-mediated endocytosis of AMPARs, where its overexpression reduces AMPAR surface levels and its deletion impairs long-term depression (LTD; Xiao et al., 2006; Davidson et al., 2009). Behaviorally, NSG3 overexpression leads to perseverative navigation in the Morris Water Maze and deficits in fear extinction (Vazdarjanova et al., 2011). Together, these studies support a shared role for NSG proteins in regulating AMPAR trafficking at excitatory synapses, with meaningful, albeit convoluted, consequences for behavior.

However, NSG functions extend beyond postsynaptic regulation. NSG1-3 proteins are among the most upregulated proteins during neuronal development (Floruta et al, 2017), where NSG1 contributes to the transcytosis of L1CAM from dendrites to axons, a process vital for axon outgrowth (Yap et al., 2008). Both NSG1 and NSG2 interact with Sortilin-1 (SORT1), though only NSG1 facilitates ADAM10/17-mediated ectodomain shedding of SORT1 (Overby et al., 2023). Moreover, NSG1 has been linked to the production of β-CTF and amyloid-β in models of amyloid precursor protein (APP) processing (Norstrom et al., 2010). NSG3 promotes proteolytic cleavage of pro-Neuregulin-1 (NRG1) by β-secretase (BACE1), generating soluble NRG1 fragments critical for neuronal signaling (Yin et al., 2015). More recently, NSG3 was shown to mediate synapse reformation following injury through interactions with the astrocyte-secreted protein SPARCL1/HEVIN (Kim et al., 2021). Collectively, these findings highlight the multifaceted and essential roles of NSG proteins in neuronal development, signaling, and plasticity.

In this study, we sought to comprehensively define the molecular landscape of NSG protein function by mapping their interactomes using a TurboID based, proximity-dependent labeling proteomics approach (Branon et al., 2018; Kim and Roux, 2016). This strategy identified over 1,000 putative protein interactors, revealing both overlapping and unique interaction profiles among NSG family members. Bioinformatic analysis confirmed known associations with Golgi, endosomal, and plasma membrane compartments, and reinforced established links to glutamatergic synapses and ADAM10-mediated proteolysis. Intriguingly, the data also pointed to potential new roles in presynaptic and inhibitory synaptic function, protein synthesis regulation, and subcellular protein sorting—particularly through interactions with multi-subunit tethering complexes such as retromer. These findings lay the groundwork for more targeted hypotheses about NSG protein function, offering both a broad systems-level view and a refined framework for dissecting the distinct biological roles of each NSG family member.

## RESULTS

### TurboID fusion constructs show normal subcellular trafficking

We mapped the protein interactome of NSG family members by designing fusion constructs linking the promiscuous biotin ligase TurboID (Branon et al., 2018) and a V5 epitope tag to the cytoplasmic domains of NSG1-3. NSG1 and NSG2 are type II transmembrane (TM) proteins with a cytoplasmic amino-termini (N-termini) (Yap et al., 2017), while NSG3 is a type I TM protein, with cytoplasmic carboxy-terminus (C-terminus) (Xiao et al., 2006). Figure 1A and 1B illustrate a diagrammatic summary of the experimental design, from transgene expression and biotinylation in human neurons, to protein purification and analysis of peptides by liquid chromatography-mass spectrometry (LC-MS). To enhance the human relevance and translatability of our findings, we expressed TurboID constructs in neurons derived from human induced pluripotent stem cells (iPSCs) containing a doxycycline-inducible Neurogenin-2 (i^3^N iPSC line; Wang et al., 2017). These induced human neurons, commonly referred to as “i^3^Neurons”, are highly scalable, reach functional maturation levels similar to rodent primary cultures, and do so on an accelerated timeline compared to directed differentiation of hPSCs (Johnson et al., 2007; Weick et al., 2011).

**Figure 1:**
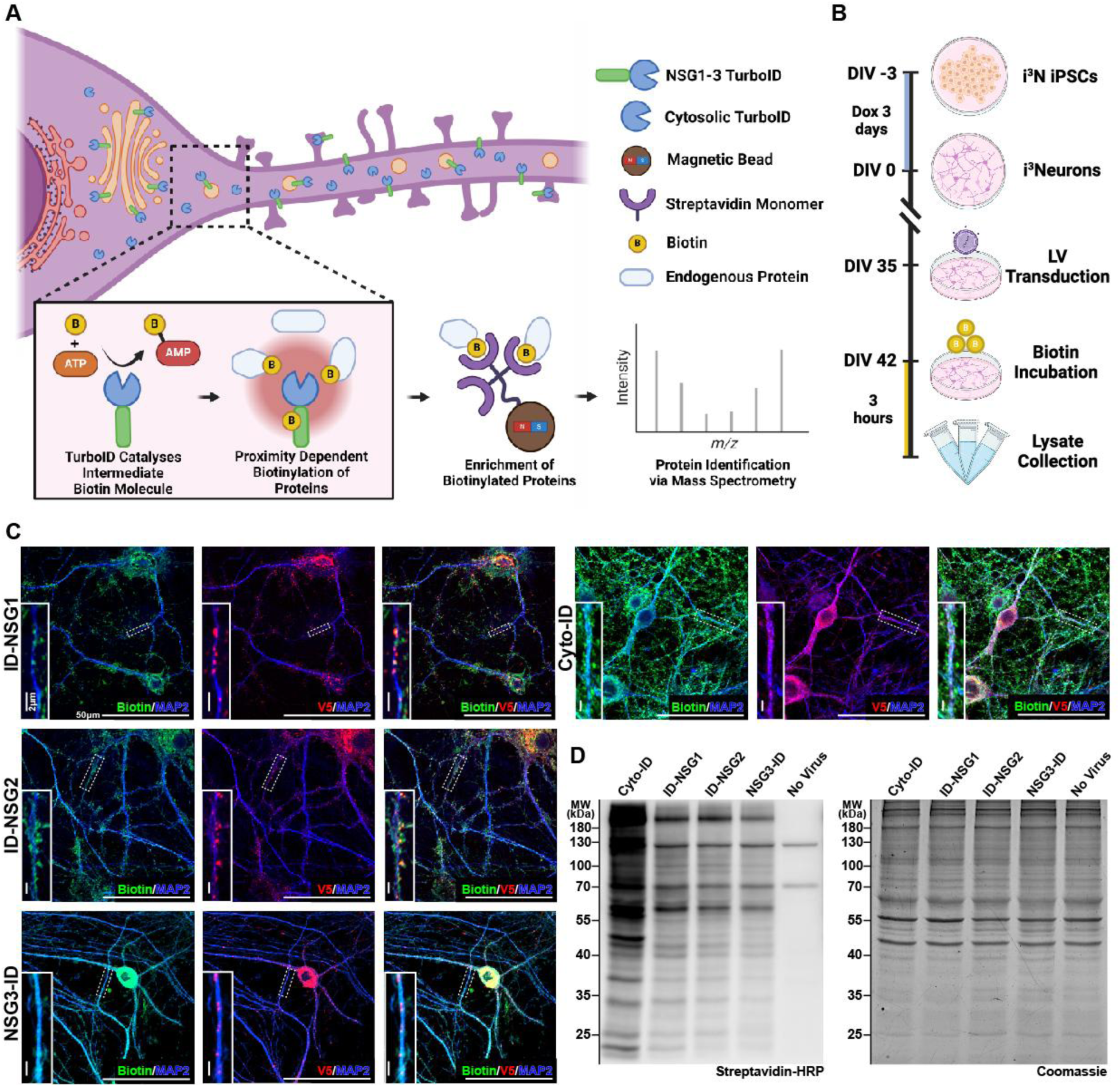
Experimental design and validation of Turbo-ID constructs. (A) Diagrammatic summary of TurboID-based methods for promiscuous biotinylation of proteins in close proximity of ID-NSG1, ID-NSG2, NSG3-ID, and cytoplasmic control (Cyto-ID). (B) Experimental timeline indicating i^3^N differentiation, lentiviral transduction of NSGX-IDs or Cyto-ID control, biotinylation, and harvesting of neuronal lysates for mass spectrometry-based analysis. (C) Immunocytochemistry (ICC) labeling of each TurboID-fusion protein demonstrates accurate subcellular targeting and biotinylation. Scale bars, 50µm and 2µm. (D) Western blot analysis displaying biotinylated proteins (left) and total protein stain (right) from whole neuronal lysates across NSGX-IDs and controls.

To control for non-specific biotinylation we used TurboID linked to a nuclear export signal (Branon et al., 2018) which localizes to the cytoplasm (Figure 1C, right panel; referred to as Cyto-ID). Proper subcellular targeting of overexpression vectors was determined via fluorescent immunolabeling of the V5 epitope tag as well as the endogenous NSG proteins in cytochemical (Kruusmägi et al., 2007; Yap et al., 2017; Chander et al., 2019). All NSG1-3:TurboID fusion constructs (hereafter collectively referred to as NSGX-ID) were expressed in a peri-nuclear pattern, consistent with Golgi localization, as well as in a punctate distribution throughout MAP2**^+^**dendrites, consistent with their presence in multiple vesicular organelles (Figure 1A; Chander et al., 2019; Yap et al., 2017, Steiner er al., 2002, Ali and Bergson, 2003; Saberan-Dejanedi e, 1998). We observed extensive colocalization of the V5 tag (red) and NSG1-3 targeting antibodies, with negligible signal for NSG1-3 (green) punctae alone, indicating proper localization of NSGX-ID constructs (Supplemental Figure 1A). Additionally, we observed robust biotin labeling across all constructs using a streptavidin-conjugated Alexa Fluor 488 fluorophore (Figure 1C, green). While a significant portion of V5 tag signal co-localized with the biotin label, we also observed biotinylated proteins that did not co-localize with V5 tag punctae (Figure 1C). This is likely due to movement of NSG1-3 themselves and/or proteins that transiently interact with, and are biotinylated by, NSGX-IDs during protein trafficking (Muthusamy et al., 2015; Yap et al., 2017). Protein biotinylation and efficient biotinylated protein enrichment was further verified via immunoblotting of neuronal cell lysates across conditions in both whole lysates and enriched protein eluates (Figure 1D and Supplemental Figure 1B, see methods).

### NSG1-2 have similar interactomes, divergent from NSG3

We performed Liquid Chromatography-Mass Spectrometry (LC-MS) analysis on neuronal cell lysates from three biological replicates, and two technical replicates per sample, across all four groups (24 samples total). Following quality control and quantitative analysis of peptide intensities, we identified a total of 2410 proteins that met the inclusion criteria (see methods) in at least one of the experimental or control groups (Supplemental Table 1). Across proteins identified in NSGX-IDs and Cyto-ID groups, abundance levels were consistent, with a median coefficient of variation (CV) ranging from 12.5% and 21.6% (Supplemental Figure 2A & 2B). Figure 2A shows hierarchical cluster analysis (HCA) along with a heat map of relative abundances of individual proteins, demonstrating a high degree of replicability across biological and technical replicates within groups, as well as notable differences between groups. Principal Component Analysis (PCA) also demonstrated tight clustering of technical and biological replicates within groups and indicates minimal variation across replicates. Interestingly, all NSGX-ID experimental groups clustered away from Cyto-ID, indicating unique interactomes compared to the cytoplasmic control. Furthermore, ID-NSG1 and ID-NSG2 showed relatively tight clustering with each other (Figure 2B, blue and red circles), with significant divergence from NSG3-ID samples (Figure 2B, gray circles) and Cyto-ID (Figure 2B, black circles).

**Figure 2:**
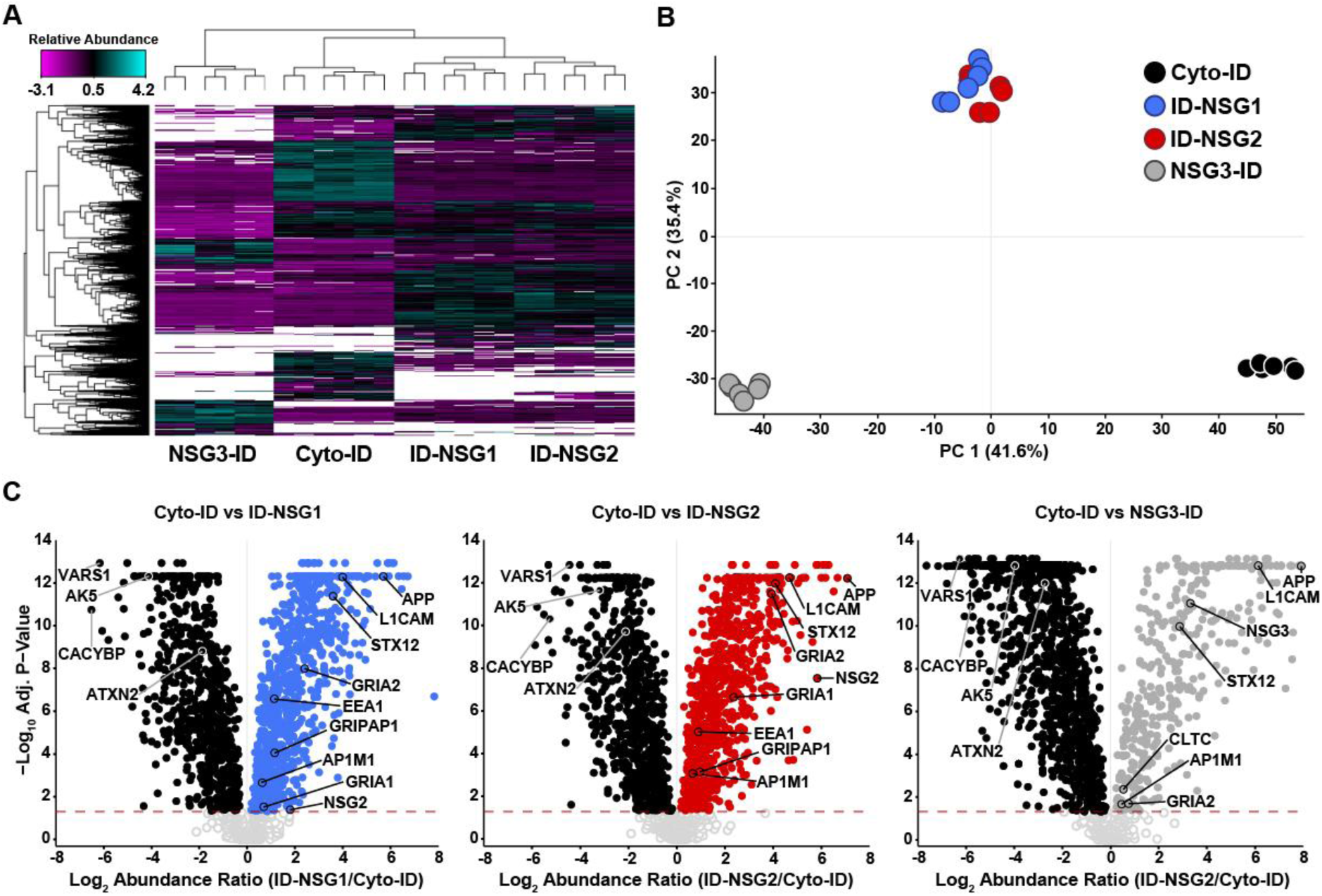
NSG1-2 have similar interactomes, divergent from NSG3. (A) Hierarchical cluster analysis of 2410 proteins identified across all four groups demonstrates highly reproducible results across 3 biological and 2 technical replicates (n=6 per group). Relative abundance z scores depicted with a pseudo-color scale. (B) Principal component analysis (PCA) scatter plot demonstrates significant differences of all NSGX-ID groups and Cyto-ID controls, as well as similarity between interactomes of ID-NSG1 and ID-NSG2. Each circle represents a technical replicate. (C) Volcano plots of significantly enriched proteins in the Cyto-ID group (filled black dots) compared with NSGX-ID groups (filled blue, red, or gray dots). Dashed line marks adjusted p-value = 0.05. Proteins known to be cytoplasmic as well as known interactors of NSG1-3 or localized to NSG1-3-containing vesicles are labeled in each group.

Given our approach, protein abundances are a correlate measurement for biotinylation levels and thereby extent of interaction. When compared to Cyto-ID controls, proteins with normalized abundance ratios > 1 (NSGX-ID/Cyto-ID) and an adjusted significance threshold of p < 0.05 in at least one of the NSGX-ID groups were considered, 947 proteins were significantly enriched in the ID-NSG1 group, 957 proteins enriched in the ID-NSG2 group, and 456 proteins in the NSG3-ID group (Supplemental Figure 2C). We found 1144 unique proteins significantly enriched across all NSGX-ID groups, of which 27.1% (310/1144) were shared across all NSGX-ID groups (Supplemental Figure 2D). The ID-NSG1 and ID-NSG2 interactomes showed the greatest overlap, exclusively sharing 48.4% (554/1144) of NSGX-ID enriched proteins, with only 6.8% (64/947) and 7.3% (70/957) of their respective interactomes diverging. In contrast, NSG3-ID had a greater degree of divergence, with 22.8% (104/456) of its interactome specific to NSG3-ID (Supplemental Figure 2D).

To visualize which specific proteins were most significantly different between groups, we generated volcano plots for each NSGX-ID/Cyto-ID comparison. Figure 2C illustrates significantly abundant proteins for each NSGX-ID (blue, red, gray dots) as well as those that were significantly enriched in the Cyto-ID group (black dots). Many cytoplasmic proteins were enriched in the Cyto-ID group including VARS1, CACYBP, ATXN2, and AK5 as expected (Filipek et al., 2002; Solaroli et al., 2009; Thul et al., 2017). Further, multiple proteins previously shown to interact with NSG family members were enriched in the expected NSGX-ID group (Muthusamy et al., 2015; Chander et al., 2019; see below). Interestingly, each NSG1-3 protein was found among the greatest significant fold increases in their respective experimental group compared to controls (i.e., ID-NSG2 biotinylated NSG2). A small subset of proteins, including NSG1, did not reach the threshold criteria for inclusion in the Cyto-ID controls and therefore no statistical comparison could be made with Cyto-ID (Supplemental Table 1). However, since NSG1 protein levels were significantly increased in ID-NSG1 compared to ID-NSG2 and NSG3-ID levels (Figure 4B), together these data are suggestive of auto-biotinylation and/or homodimerization with endogenous NSG proteins (see below).

### Pathway analyses reveal potential novel roles of NSG proteins

In order to shed insight into the molecular pathways enriched in the NSG1-3 interactome, we first performed a gene set enrichment analysis (GSEA) using the Gene Ontology (GO) and Kyoto Encyclopedia of Genes and Genomes (KEGG) databases on the list of 1144. Across NSG family interactors, significantly enriched GO categories within the Biological Process and Molecular Function clades included multiple predicted processes relating to synaptic signaling and organization, as well as involvement in transmembrane transport of ions (Figure 3A & 3C), consistent with the role of NSGs in regulating glutamate receptors at synapses. The Cellular Component clade revealed known organelle localization such as “plasma membrane”, “synaptic membrane”, and “early endosome” as significantly enriched (Figure 3B). KEGG pathway enrichment revealed categories such as “endocytosis”, “glutamatergic synapse”, and “long-term potentiation” as significantly enriched (Figure 3D). Together, these data provide further confidence in the fidelity of the experimental system by corroborating established roles of NSG family members. Interestingly, GSEA of all GO clades and KEGG pathways returned additional categories not previously associated with NSG1-3 protein function. Many enriched terms and pathways pertained to regulation of GABAergic synapse function such as “synaptic transmission, GABAergic” and “GABAergic synapse”, among others (Figure 3A-D, orange text). We also noted terms and pathways representing presynaptic functions such as “presynaptic membrane”, “SNARE interactions in vesicular transport”, and “axon guidance” (Figure 3A-D, purple text). Both of these categories are intriguing as NSG proteins have not been investigated relative to GABAergic function, and have been explicitly omitted from presynaptic analysis due to their apparent restriction to somatodendritic compartments (Chander et al. 2019; Yap et al., 2017). Finally, the appearance of endoplasmic reticulum (ER) and ribosome categories is intriguing as NSGs have not previously been associated with protein translation (Figure 3A-D, turquoise text). The complete list of GO and KEGG terms can be found in Supplemental Table 2.

**Figure 3:**
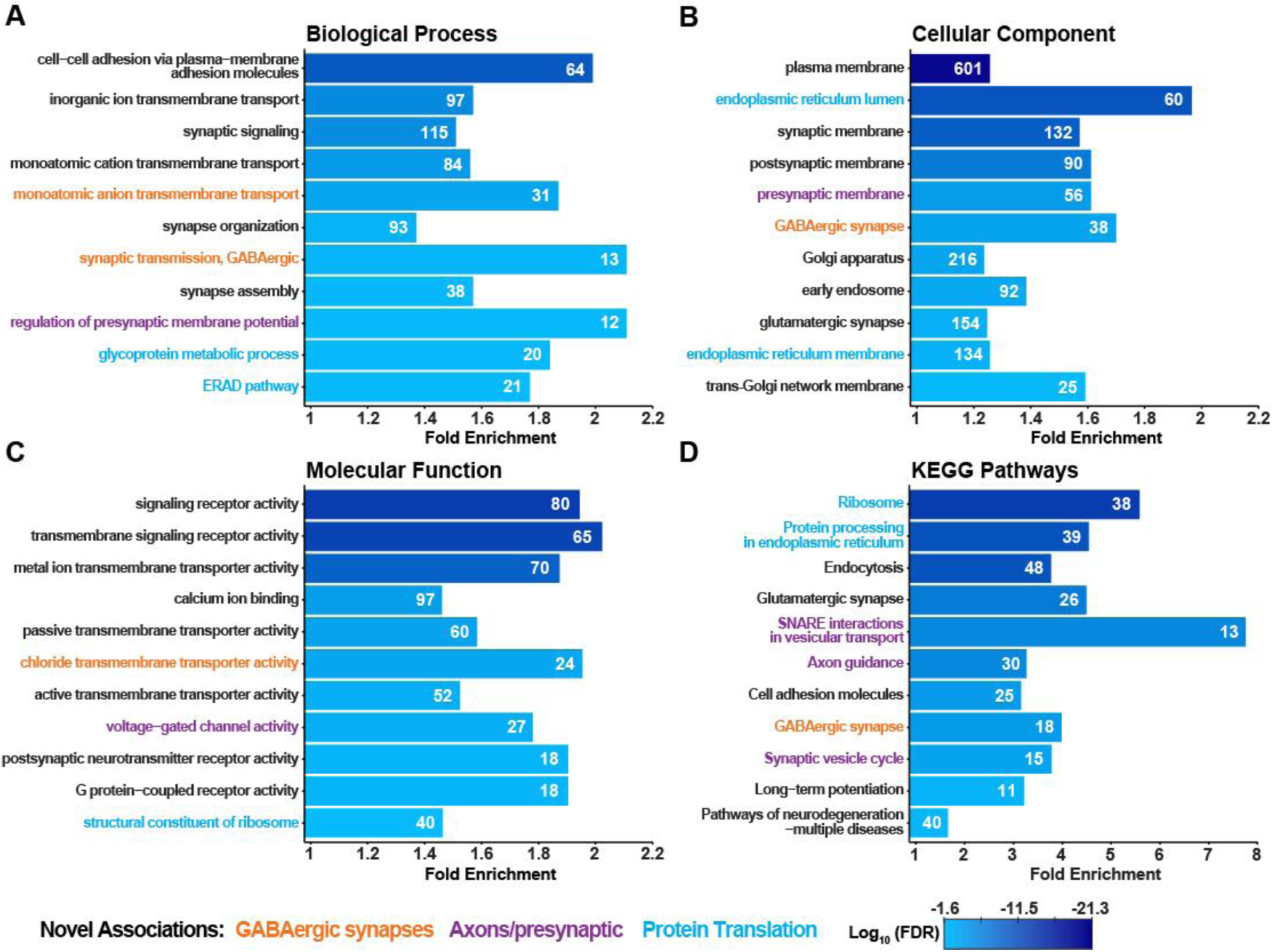
GO and KEGG analysis of NSG1-3 interactomes reveals known and novel pathway involvement. (A-C) Gene Ontology and (D) KEGG Pathways GSEA of 1144 proteins enriched across all NSGX-ID groups demonstrates predicted involvement in glutamatergic synaptic function, ion and G-protein coupled receptor activity, as well as endosomal trafficking. GO categories also identified enrichment of novel categories such as GABAergic synapse function (orange), presynaptic function (purple), as well as protein translation in the endoplasmic reticulum (turquoise). KEGG Pathways also found NSG family interactome genes representative of pathways of multiple neurodegenerative processes. Bar graphs shows the fold enrichment for each term or pathway, numbers inside bars indicate the number of genes driving enrichment. All categories have a log_10_ (FDR) < 0.05, represented by pseudo-colored scale.

In an attempt to identify unique functions of individual NSG family members, we identified significantly enriched GO & KEGG categories for individual NSGX-ID groups (Supplemental Table 2). Segregation of GO and KEGG categories was readily performed for NSG3, which demonstrated that NSG3 was primary driver of pathways pertaining to the ER and protein translation (Supplemental Figure 3A-D, turquoise text). Unique pathway enrichment for ID-NSG1 and ID-NSG2 was more difficult due to their significantly overlapping interactomes, which appeared to be the primary drivers of synaptic and endosomal GO/KEGG pathway enrichment (Supplemental Figure 3A-D, green text). Despite this overlap, uniquely enriched categories for ID-NSG1 included “Huntington’s disease”, “Alzheimer’s disease”, as well as “Prion disease”, while those enriched in the ID-NSG2 group included multiple ‘SNARE’ complex categories and “voltage-gated potassium channel activity”. Despite the fact that both NSG1 and NSG2 both bind the AMPAR subunits it was notable that “glutamate receptor binding” and “AMPAR receptor complex” were enriched in the ID-NSG2 group (Supplemental Figure 3B & 3C).

To further attempt to segregate molecular functions of individual NSG family members, we curated lists of proteins that were preferentially enriched in a single group. GO and KEGG GSEA of these severely truncated lists (ID-NSG1 (142), ID-NSG2 (103), and NSG3-ID (251); Supplemental Table 1) returned few results (Supplemental Table 3). Thus, we generated protein-protein interaction (PPI) networks using the STRING database which clusters and annotates proteins based on predominantly known and predicted functions of the grouped PPI (Supplemental Figure 4A-C; Szklarczyk et al., 2023, 2021, 2019). While clustering of proteins in the ID-NSG1 and ID-NSG2 groups was less apparent, ID-NSG1 had the largest specific association with adherens junction proteins, as well as annotations related to protein trafficking such as Retromer complex, TRAPP complex, and RHOQ GTPase cycle (Supplemental Figure 4A). Pathways specifically associated with NSG2 included RHOD GTPase cycling, activation of AMPA receptors, and ER-Golgi transport (Supplemental Figure 4B). Here again the NSG3-ID group had clustering of proteins related to protein processing in the ER and mRNA processing, with additional groups related to plasma lipoprotein particles and presynaptic functions (Supplemental Figure 4C). However, many NSGX-ID-specific proteins were ungrouped either indicating a significant overlap of NSG function within pathways or incomplete associations in the database.

### Proteomic data support previously established NSG1-3 protein interactions

For identification of potentially significant NSG interacting partners, we used abundance ratios of NSGX-ID normalized to Cyto-ID controls, which also allowed us to look for significant preferential enrichment by comparing abundance ratios across NSGX-ID groups. In cases where no biotinylation occurred in the Cyto-ID group, we assigned a pseudocount (PC) value of “1” to replace the missing abundance value. This resulted in very large ratios (typically orders of magnitude) which could not be analyzed statistically, but may represent some of the more critical NSG-interacting partners within subcellular domains that exclude Cyto-ID. Previous studies identified a number of NSG binding proteins such as Adaptor Protein mu subunits (e.g. AP1M1) subunits, Clathrin (CLTC), L1 cell adhesion molecule (L1CAM), pro-Neuregulin-1 (NRG1), PIKFYVE, as well as Syntaxin-12 (STX12; Chander et al., 2019; Ha et al., 2012; Muthusamy et al., 2012; Steiner et al., 2005, Qi et al., 2023; Xiao et al., 2006; Yin et al., 2015). Figure 4A demonstrates significant biotinylation ratios for multiple NSG family members over Cyto-ID (Cyto-ID control levels indicated by dashed line at “1”) for AP1M1, CLTC, L1CAM, NRG1, PIKFYVE, and STX12. However, many known interactors were not identified in the current dataset (see discussion).

**Figure 4:**
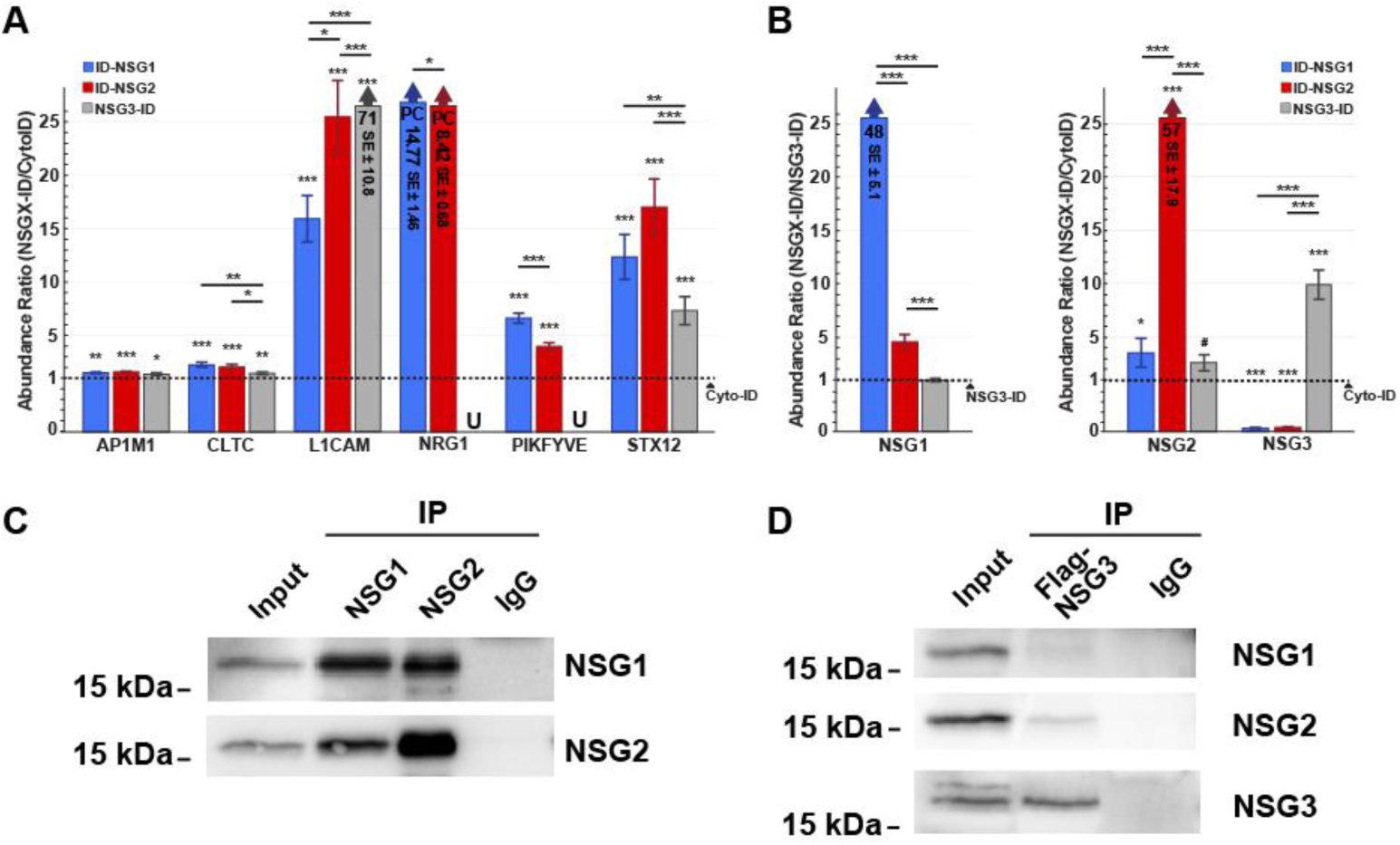
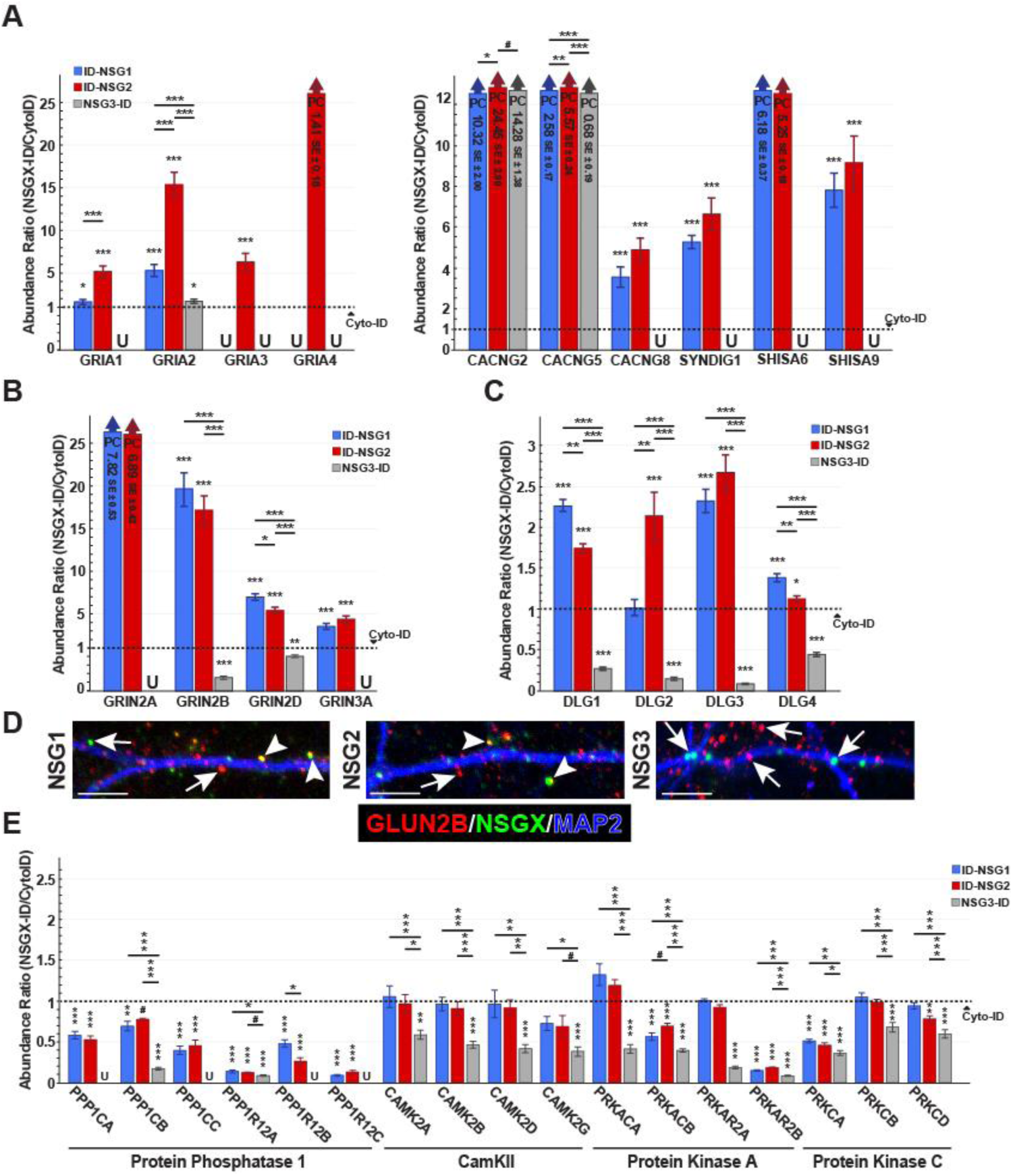
Interactome reveals heteromultimerization between NSG proteins. Abundance ratios of NSGX-ID groups compared to Cyto-ID control, unless stated, (dashed line at “1”) show: (A) previously established interactors of NSG proteins are significantly enriched (See below for annotations). (B) Auto-biotinylation of NSGX-ID groups. Left panel shows NSGX-ID groups compared to ID-NSG3 for NSG1 protein as it was not identified in Cyto-ID controls. Right panel shows NSGX-ID compared to Cyto-ID control for NSG2 and NSG3 proteins. (C) Whole mouse brain Co-Immunoprecipitation of endogenous NSG1 and NSG2 revealed physical interactions between both proteins. IgG control antibody was not able to precipitate either NSG1 or NSG2. (D) Overexpression of a FLAG-tagged NSG3 protein in human i3Ns revealed successful Co-IP with the anti-Flag antibody (bottom blot), but very modest interactions with NSG1 and NSG2 (upper two blots), while control IgG could not precipitate any NSG family member. For all bar graphs, “PC” labeled bars show ratio and standard error (SE) in millions. Non-PC bars that have a large abundance ratio were restricted, ratio and SE stated inside bar. *# p < 0.1, *p < 0.05, **p < 0.01, ***p < 0.001*.

### NSG1-3 differentially bind each other and form heteromultimeric complexes

In addition to known interacting proteins, NSGX-ID constructs biotinylated their corresponding NSG protein to a significantly greater extent than Cyto-ID (Figure 4B; Supplemental Table 1), suggesting either self-biotinylation or homomultimerization with endogenous NSG1-3 proteins. Interestingly, we also found that NSGX-ID constructs significantly biotinylated *other* NSG family members in addition to themselves. This was particularly evident for NSG1-ID and NSG2-ID, which robustly biotinylated both NSG1 and NSG2 to a significantly greater extent than Cyto-ID (Figure 4B). In contrast, ID-NSG1 and ID-NSG2 biotinylated NSG3 significantly *less* than Cyto-ID (Figure 4B, right, blue/red bars). However, NSG3-ID did biotinylate NSG1 and NSG2 to levels near significance compared to Cyto-ID, indicating that the relationship between NSG3 and the other two proteins may be complex, and that the TurboID construct itself may promote or inhibit this interaction due to steric effects or expression levels (see discussion). To determine whether NSG proteins potentially form heteromeric complexes we used co-immunoprecipitation (Co-IP) of each NSG family member and probed for the presence of other family members. Figure 4C shows that endogenous NSG1 and NSG2 immunoprecipitated from mouse brain samples were able to reciprocally precipitate each other (lanes 2-3), while a control IgG antibody didn’t precipitate either protein. While we were unable to detect endogenous NSG3 (not shown), we pulled down overexpressed NSG3 linked to a FLAG tag (see methods; Ali and Bergson, 2003) from human i3Ns. Precipitation of FLAG-NSG3 revealed robust auto-precipitation (Figure 4D, lower blot), but very weak bands for NSG1 and NSG2 (Figure 4D, upper blots). IgG control showed no specific precipitation product (Figure 4D, all blots). Together, these data suggest that NSG1 and NSG2 form heteromeric complexes, and the possibility of a small percentage of complexes that also contain NSG3.

### Interactome data support established and novel interactions with excitatory postsynaptic proteins

As mentioned, glutamatergic synaptic transmission was one of the most significantly enriched GO and KEGG pathways (Figure 3). All NSG family members are well-characterized for their regulated trafficking of AMPA receptor subunits GLUA1 (GRIA1) and GLUA2 (GRIA2) (Alberi et al., 2005; Davidson et al., 2009; Chander et al., 2019), which were both significantly abundant in ID-NSG1 and ID-NSG2 groups (Figure 5A). Interestingly, subunits GLUA3 (GRIA3) and GLUA4 (GRIA4) were also enriched, but only in the ID-NSG2 group (Figure 5A). NSG2 also showed significantly greater biotinylation of all AMPAR subunits compared with ID-NSG1 and NSG3-ID (Figures 5A), while GLUA2 was significantly abundant for NSG3-ID. AMPARs associate with a number of auxiliary subunits both during trafficking and at the plasma membrane, including TARPs, Cornichons (CNIH), SynDIG1, and Shisa6,9 (CKAMP44 and 52; Farrow et al., 2015) (Schwenk et al., 2012). Of these, we identified multiple members of the TARP family (CACNG2, CACNG5, and CACNG8), both Shisa6 and 9 (CKAMP44 and 52), and SynDIG1 as significantly biotinylated in NSGX-ID groups (Figure 5A, right). Together with previous immunoprecipitation data (Alberi et al., 2005; Chander et al., 2019), these results suggest that NSG1-3 bind mature AMPARs during secretory or endosomal trafficking, and/or at the plasma membrane.

**Figure 5:**
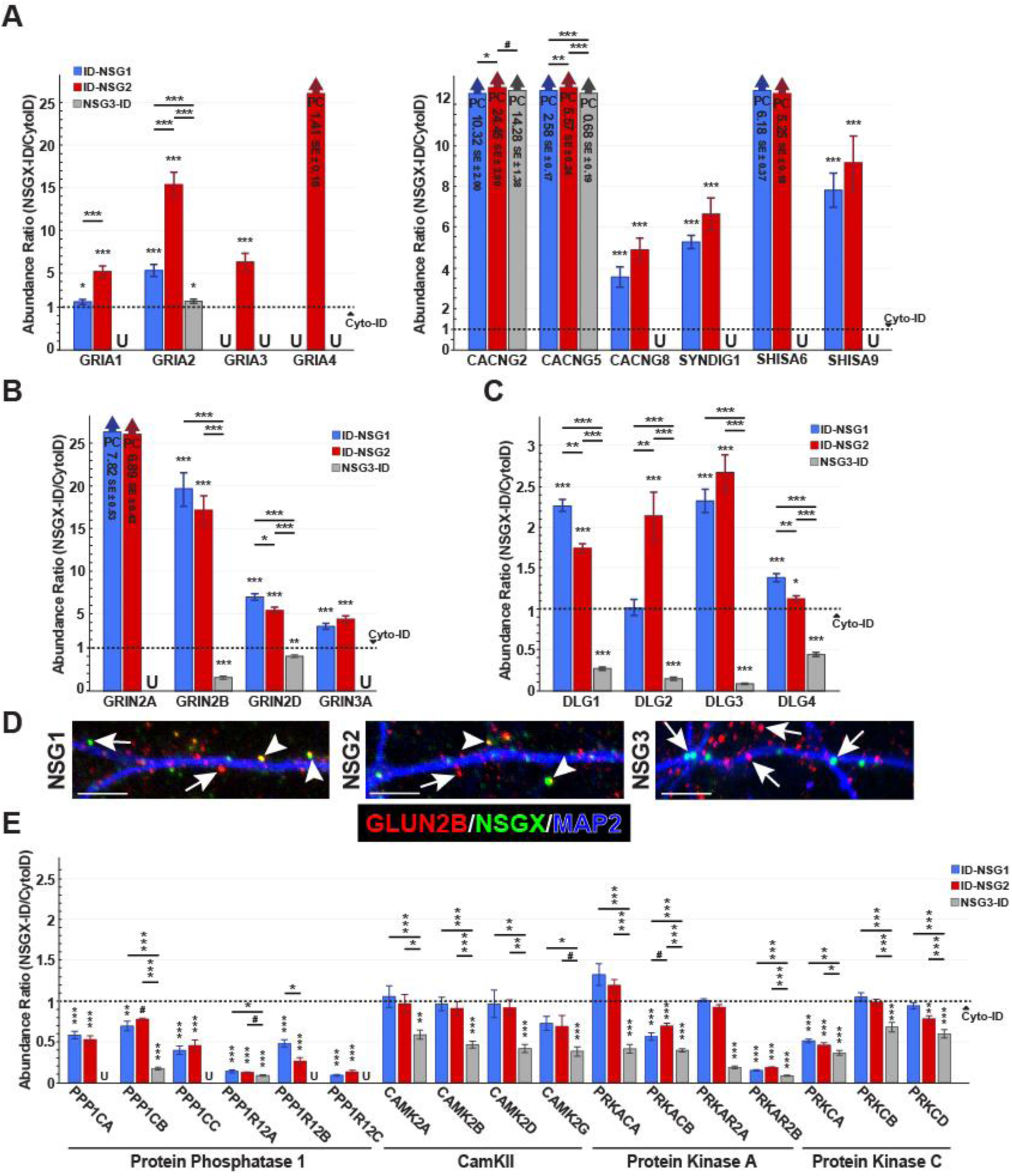
Interactome data support established and novel interactions with excitatory postsynaptic proteins. Abundance ratios of NSGX-ID groups compared to Cyto-ID control (dashed line at “1”) show: (A) all AMPA receptor subunits (GLUA1-4; left) and several AMPA receptor auxiliary proteins (right). (B) Several NMDAR subunits and (C) Postsynaptic density (PSD) scaffolding proteins belonging to the MAGUK family. (D) ICC colocalization of endogenous NSG1-3 and GluN2B in cultured mouse neurons. Scale bars are 10µm. (E) Subunits of major kinases and phosphatases involved in synaptic function. For all bar graphs, “U” denotes undetected proteins in the corresponding NSGX-ID group. “PC” labeled bars show ratio and standard error (SE) in millions. Non-PC bars that have a large abundance ratio were restricted, ratio and SE stated inside bar. *# p < 0.1, *p < 0.05, **p < 0.01, ***p < 0.001*.

In addition to AMPARs and auxiliary subunits, we were surprised to find multiple subunits of NMDA receptors (NMDARs) significantly abundant in ID-NSG1 and ID-NSG2 groups (Figure 5B). However, while GRIN2A, 2B, and 3A were identified, the GRIN1 subunit was conspicuously absent from all groups, including Cyto-ID. As homomers of GRIN1 or GRIN2 subunits are not trafficked to the plasma membrane (McIlhinnet et al., 1998), this suggests either a unique type of interaction with receptor subunits prior to assembly, steric hinderance of NSG-ID proteins by GRIN2 subunits, or technical limitations in the identification of GRIN1 using these methods. However, we found robust ICC co-localization was between both NSG1/2 and the GRIN2B subunit (Figure 5D), suggesting a possible indirect role of NSG1-2 in mediating synaptic NMDAR trafficking (Washbourne et at., 2004). ID-NSG1 and ID-NSG2 also significantly biotinylated post-synaptic scaffolding proteins of the Disks Large/MAGUK family (DLG1-5; Figure 5C) and their associated proteins (DLGAP-1-4; Supplemental Figure 5A; Bai et al., 2022; Sheng and Hoogenraad et al., 2007), with NSG3-ID uniquely biotinylating DLGAP4 greater than Cyto-ID controls (Supplemental Figure 5A). Additional excitatory synaptic proteins biotinylated by NSGX-IDs included trans-synaptic adhesion proteins including the human-specific NLGN4X (Supplemental Figure 5C), signaling molecules such as NOS1 (Supplemental Figure 5B), the metabotropic glutamate receptor 5 (GRM5), and other scaffolding proteins (Supplemental Figure 5). In contrast, nearly all synaptic kinases and phosphatases including PP1, CaMKII, PKA and PKC were either significantly less biotinylated by NSGX-ID proteins, or not different from Cyto-ID controls (Figure 5E). Together these data support a broader role of NSG1-3 within excitatory post-synaptic regions rather than simply as AMPAR-trafficking proteins.

### Potentially novel roles for NSG1 and NSG2 in presynaptic and inhibitory transmission

Nearly all studies on NSG family members to date have found them restricted to the somatodendritic compartment (Yap et al., 2017; Chander et al., 2019). However, KEGG and GO enrichment analysis found significant enrichment for pathways such as axon guidance and presynaptic membrane (Figure 3B & D, Supplemental Figure 3, purple text), indicating a potential role for NSGs in the axon or bouton. Within these pathways, we identified a number of presynaptic proteins biotinylated by NSGX-ID to a significantly greater extent than Cyto-ID. These include scaffolding proteins (PCLO, BSN, CASK, RIMS1), SNARE complex proteins (SNAP25, VAMP2, STX1B, SYT7/11), voltage-gated calcium channels (CACNA1B, CACNA1E), trans-synaptic adhesion proteins (NLGN3/4X), and the Thrombospondin receptor CACNA2D1 (Figure 6A; Rizalar el al., 2010; Chia et al., 2013). These data prompted us to revisit the potential pre-synaptic localization of NSG1-3, where ICC analysis revealed a small, but significant proportion of Synaptophysin-1**^+^**boutons co-localized with each family member (Figure 6C). Together, these data indicate that NSG family members may play an important role in the trafficking of proteins involved presynaptic assembly, maintenance, and vesicle release but how and where NSGs interact with these proteins remains to be studied (see discussion).

**Figure 6:**
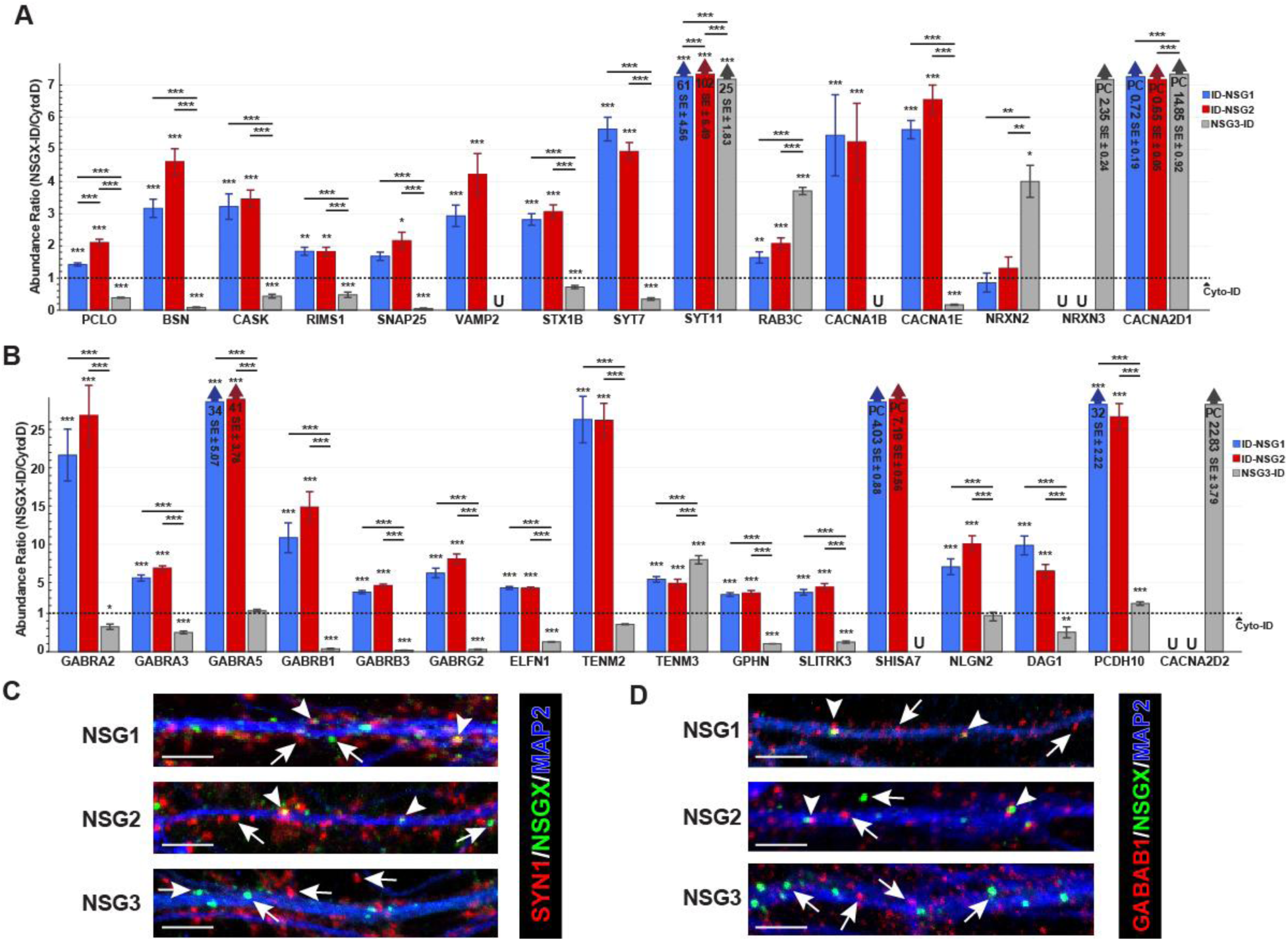
Potentially novel role for NSGs in presynaptic boutons, inhibitory synapses. Abundance ratios of NSGX-ID groups compared to Cyto-ID control (dashed line at “1”) show: (A) presynaptic markers involved in different mechanisms such as scaffolding, transsynaptic adhesion, synaptic vesicle release, and ion transport are significantly biotinylated across NSGX-IDs. (B) GABAergic synapse markers including various GABA_A_ Receptor subunits, scaffolding proteins, and transsynaptic adhesion proteins are primarily enriched in ID-NSG1 and ID-NSG2. Confocal images of (C) presynaptic marker SYN1 or (D) GABA_A_ Receptor subunit β2 and endogenous NSG1 or NSG2 in mouse cultured neurons. For NSG3 colocalization, we transfected NSG3-mNGreen into mouse cultured neurons. A portion of NSG1 and 2 punctae colocalized (arrowheads) with SYN1 or β2. Full arrows point non-colocalized signal. For all bar graphs, “U” denotes undetected proteins in the corresponding NSGX-ID group. “PC” labeled bars show ratio and standard error (SE) in millions. Non-PC bars that have a large abundance ratio were restricted, ratio and SE stated inside bar. *# p < 0.1, *p < 0.05, **p < 0.01, ***p < 0.001*.

KEGG pathway and GO enrichment analysis also identified proteins associated with inhibitory neurotransmission at GABAergic synapses (Figure 3), which has not previously been reported. The post-synaptic ionotropic GABA_A_ Receptor is comprised of five subunits with a stoichiometry of two alpha, two beta and one gamma, or less commonly a delta subunit (Krueger-Burg et al., 2025). Subunits encoding α2, α3, α5, β1, β3, γ2, and γ3 were all significantly biotinylated by ID-NSG1 and ID-NSG2 compared to Cyto-ID controls, while NSG3-ID showed significantly less biotinylation for all subunits (Figure 6B). Similar to NMDARs, we were surprised to find that the α1 subunit was not identified in any group including Cyto-ID, given the fact that a large majority of native GABA_A_ receptors contain α1 (Washbourne et al., 2004; Lau and Zukin et al., 2007). Additional proteins important for GABAergic synapse assembly and function were identified, including the post-synaptic scaffold proteins Gephyrin and Slitrk3, as well as the trans-synaptic adhesion protein Neuroligin-2 (NLGN2; Figure 6B). Using subunit β1 (GABRB2) as proxy for GABA_A_ receptors we looked for ICC colocalization with NSG1-3, we found a portion of NSG1 and NSG2 punctae colocalizing with GABRB signal. In contrast less localization was observed with NSG3. Our data indicate that NSG modulation of synaptic plasticity via receptor trafficking may extend beyond excitatory synaptic function.

### NSGs interact with markers of multiple subcellular compartments

NSGs regulate the trafficking of protein substrates within multiple intracellular vesicular compartments (Steiner et al., 2002 & 2005, Xiao et al., 2006, Yap et al., 2008, Muthusamy et al., 2012, Chander et al., 2019; Yap et al., 2017). However, the full panoply of subcellular locations and precise mechanisms for NSG-mediated trafficking remain unknown. As NSG1 and NSG2 were originally identified in Golgi apparatus (Saberan-Djoneidi et al.,1995, 1998), multiple Golgi markers were identified as expected (Figure 7A). Interestingly, NSGs appeared to be specifically enriched in COPII-containing vesicles, as COPII proteins were biotinylated to a greater degree than COPI proteins (Supplemental Figure 6A; Dell’Angelica and Bonifacino, 2019).

**Figure 7:**
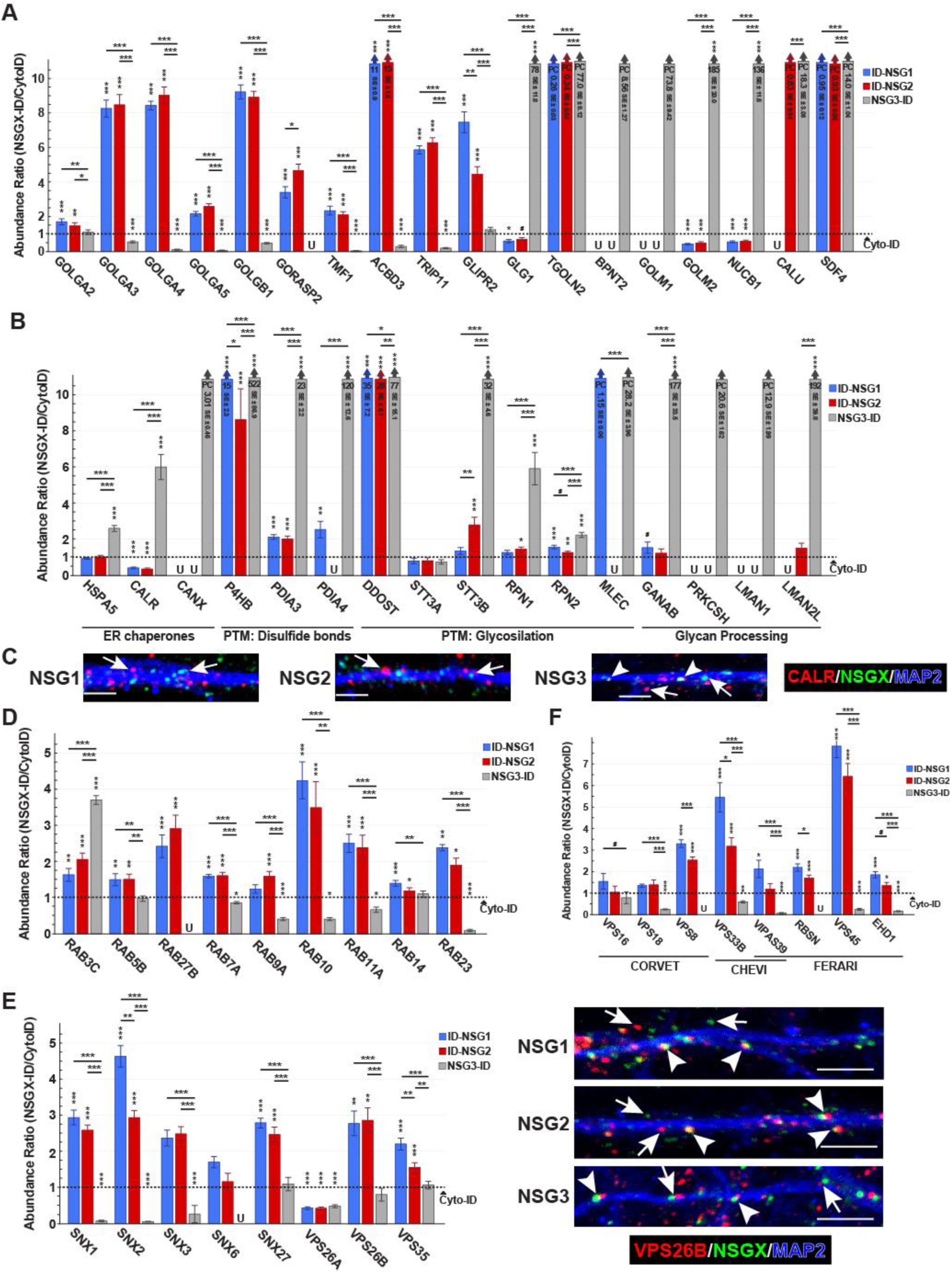
NSGs occupy multiple subcellular compartments. (A) Abundance ratios of NSGX-ID groups compared to Cyto-ID control (dashed line at “1”) show various Golgi apparatus proteins involved in organelle tethering, vesicle recognition and sorting, and protein processing with subsets being preferentially enriched by different NSGX-IDs. (B) ER proteins involved in several PTMs where NSG3-ID shows significantly greater biotinylation compared to Cyto-ID and ID-NSG1/2. (C) Confocal images of labeled endogenous NSG1 or NSG2 and the ER marker Calreticulin (CALR) in cultured neurons. For NSG3 colocalization, we transfected NSG3-mNGreen into mouse cultured neurons. A portion of NSG3 punctae colocalized (arrowheads) with CALR. Full arrows point non-colocalized signal. Scale bars are 10µm. (D) Relative biotinylation ratios of various Rab proteins involved in vesicle fusion and trafficking. (E) Left, NSG1-ID and NSG2-ID displayed abundant biotinylation of multiple components of the Retromer complex. Right, NSG1 and NSG2 also showed abundant co-localization (arrowheads) with the neuron-specific VPS26B. Scale bars are 10µm. (F) Multiunit tethering complex subunits involved in vesicle recycling are enriched in NSG1/2-ID groups. For all bar graphs, “U” denotes undetected proteins in the corresponding NSGX-ID group. “PC” labeled bars show ratio and standard error (SE) in millions. Non-PC bars that have a large abundance ratio were restricted, ratio and SE stated inside bar. *# p < 0.1, *p < 0.05, **p < 0.01, ***p < 0.001*.

Intriguingly, there was significant enrichment of ER related GO and KEGG terms, which was primarily driven by the NSG3-ID interactome (Figure 3). A detailed inspection of these pathways included well known ER proteins such as BiP (HSPA5), Calreticulin (CALR), and Calnexin (CANX) (Hebert and Molinari, 2007), all of which were only significantly biotinylated by NSG3-ID (Figure 7B). Moreover, we identified several proteins involved in post-translational modifications (PTMs) biotinylated to a greater by NSG3-ID. These included members of the PDI family which are involved in disulfide bond formation (Figure 7B; Shergalis and Neamati, 2016), as well as subunits of the OST complex known to be involved in initial protein glycosylation (Figure 7B; Hebert and Molinari, 2007; Ramirez et al., 2019). Finally, proteins involved in glycan processing (GANAB and PRKCSH) and trafficking of glycosylated proteins (LMAN1 and LMAN2L) were also only significantly biotinylated by NS3-ID (Figure 7B; Helenius and Aebi, 2004). Related, NSG3-ID also biotinylated proteins involved with RNA processing, including Ribosomal proteins and several proteins involved in RNA splicing (Supplemental Figure 6D; Jin et al., 2020; Angarola and Anczukow, 2021), potentially shedding light into a novel role of NSG3 in regulation of gene expression as this has been reported recently in a mouse pancreatic cancer model (Xia et al., 2021). Furthermore, we found a greater degree of co-localization of NSG3 with the ER marker Calretinin (Figure 7C, arrowhead) compared with NSG1 and NSG2, which showed almost no co-localization (Figure 7C, arrows). Together, these suggest a novel role for NSG3 in the initial stages of post-translational processing of newly-synthesized proteins.

To demonstrate specificity of NSG1-3 subcellular localization, we did not find enrichment of NSGX-ID biotinylated proteins that localize specifically to either peroxisomes or mitochondria. Using the MitoCarta 3.0 and PeroxisomeDB 2.0 databases we identified 48 and 5 proteins, respectively, that were shared between NSGX-ID interactomes and these databases. Of the 48 proteins, 13 are annotated as mitochondria specific and reside within the mitochondria inner membrane or matrix (Rath et al., 2020). Among the 13 proteins, 5 are endogenously biotinylated proteins MCCC1-2, PCCA, PCCB and PC, while the remaining 8 have known interactions or show STRING database correlations to these endogenously biotinylated proteins (Supplemental Figure 6B; Waldrop et al., 2012). Of the 5 shared proteins with the PeroxisomeDB 2.0, none were exclusive to peroxisome localization (Supplemental Figure 6C). Finally, we did not observe co-localization of any NSG1-3 protein with the mitochondrial marker TOMM20, nor the peroxisome marker PEX1 (Supplemental Figure 6B and 6C, right).

Rab GTPases are used for identification of specific endolysosomal compartments (Hoogenraad et al., 2010; Yap et al., 2017; Zhen and Stenmark, 2024). As expected, Early Endosome (EE) markers Rab5B and EEA1 along with associated proteins RABGEF1 and RABEP1 were enriched in ID-NSG1 and ID-NSG2 groups, although it was surprising that NSG3-ID was not different from Cyto-ID (Figure 7D, Supplemental Figure 6E). Recycling endosome (RE) markers Rab11A, RAB11FIP2/5, along with the late endosome (LE) markers Rab7A and Rab9A were also enriched (Figure 7D, Supplemental Figure 6E). We also identified several other vesicular markers that may indicate novel functions for NSGs. We identified significant biotinylation of Rab3C, Rab10, and Rab27B, in all NSGX-IDs except for Rab27b, which was not detected by NSG3-ID (Figure 7D). These Rabs are involved in anterograde trafficking of cargo through axonal compartments (Szodorai et al., 2009, Arimura et al., 2009, Deng et al., 2014) and Rab3C and Rab27B play a role in neurotransmitter release (Pavlos et al., 2010, Fischer von Mollard et al., 1994). Additionally, Rab14 and Rab23 were significantly biotinylated by ID-NSG1 and ID-NSG2, and are implicated in Golgi to RE trafficking (Evans et al., 2003, Junutula et al., 2004, Proikas-Cezanne et al., 2006) (Figure 7D), suggesting a novel role for NSG1 and NSG2 in long-distance protein recycling.

Previous studies have provided little insight into the specificity of NSG-mediated trafficking based on interactions with known sorting complexes including ESCRT, Commander, and Retromer (Verges et al., 2004, McNally et al., 2017, Cullen & Steinberg, 2018). ESCRT can be divided into four different complexes (ESCRT0-III), and is involved in protein degradation while both Commander and Retromer are involved in cargo recycling (Cullen and Steinberg, 2018). Only ESCRT-0 subunits and the TSG101 subunit from the ESCRT-I complex were found to be significantly enriched by ID-NSG1 and ID-NSG2 (Supplemental Figure 6F). We noted predominant NSG1 and NSG2 specificity for Retromer associated proteins compared to Commander complex proteins. Out of a total of seven proteins that make up the Retromer complex proper, five were significantly enriched in ID-NSG1 and ID-NSG2 groups (Figure 7E), along with a number of Retromer-associated proteins such as Rab7A, SNX3, and TBC1D5 (Figure 7D and E). Interestingly, the neuron-specific isoform of VPS26B was identified, while the ubiquitously expressed VPS26A isoform was significantly biotinylated by Cyto-ID compared to NSGX-ID (Figure 7E). In contrast, out of the sixteen Commander complex proteins (Yong et al., 2023), COMMD8 was biotinylated by Cyto-ID control alone while none were identified by any NSGX-ID group (Supplemental Table 1). To validate a potential NSG-Retromer interaction we observed significant co-localization between VPS26B and both NSG1 and NSG2 (Figure 7E, right-top panels, arrowheads). Although NSG3-ID did not significantly biotinylated Retromer proteins (Figure 7E), we observed NSG3 in close apposition to VPS26B, although NSG3 punctae were typically more adjacent to VPS26B compared with NSG1 and NSG2 (Figure 7E, right-bottom panel, arrows).

In addition to sorting complexes, multiunit tethering complexes (MTCs) modulate proper vesicle targeting and fusion (Spang, 2016, Dubuke and Munson, 2016, van der Beek et al., 2019). We identified several subunits of various MTCs, including two core subunits of the HOPS/CORVET complex and the CORVET-specific subunit VPS8 (van der Kant et al., 2015), the latter significantly enriched in ID-NSG1 and ID-NSG2 (Figure 7F). Notably, all CHEVI and nearly all FERARI subunits, a recently identified RE MTCs in mammals (van der Kant et al., 2015, Solinger et al., 2020), were significantly enriched for both or at least one NSG (Figure 7F). Both Rab5 and Rab11, the Rab GTPase counterparts of these MTCs, follow the same enrichment trend (Figure 7D).

Additionally, we also detected significant enrichment of subunits across NSGX-IDs from MTCs involved in secretory and ER-Golgi anterograde trafficking, including the Exocyst and TRAPP complexes (Wang and Hsu, 2006, Hall et al., 2024, Supplemental Figure 6G). In contrast, we failed to detect LE HOPS-specific subunits (van der Kant et al., 2015) or enrichment of retrograde trafficking MTC subunits, such as GARP or COG (Liewen et al., 2005, Ungar et al., 2002), consistent with the increased number of COPII subunits enriched for ID-NSG1 and ID-NSG2 compared to COPI subunits. These findings further support the idea that NSG1 and NSG2 are playing an important role in trafficking vesicle and/or cargo throughout the endolysosomal membrane network.

### A potentially expanded role for NSGs in regulating proteolysis of Alzheimer’s disease-related proteins

Finally, NSG family members regulate proteolysis of several disease-related proteins (Gutwein et al., 2002; Litterst et al., 2007; Parvathy et al., 1999; Norstrom 2010, Overby et al., 2023; Yin et al., 2015) that are cleaved by the “sheddase” <u>A</u> <u>D</u>isintegrin <u>A</u>nd <u>M</u>etalloproteinase domain-containing protein 10 (ADAM10). As there is no GO or KEGG pathway specifically for ADAM10 substrates, we determined whether the 90 verified mouse ADAM10 substrates (Kuhn et al., 2016) that could be mapped to the human genome were enriched in NSGX-ID groups (see methods). Thirty-four of the 90 ADAM10 substrates were significantly enriched in NSGX-ID groups compared with Cyto-ID (Supplemental Table 1) corresponding to 37% of the ADAM10 substrate list. Thirty ADAM10 substrates were biotinylated by both ID-NSG1 and ID-NSG2, which was statistically significant via two-sided Fisher’s exact test (Figure 8A). One protein was shared between ID-NSG2 and NSG3-ID, ten were shared between all groups, while only three ADAM10 substrates were solely biotinylated by NSG3-ID (Figure 8A), which was not statistically significant. A subset of ADAM10 substrates is depicted in Figure 8B, omitting duplicates included elsewhere (e.g., LINGO1, NLGN1, 3, and 4X).

**Figure 8:**
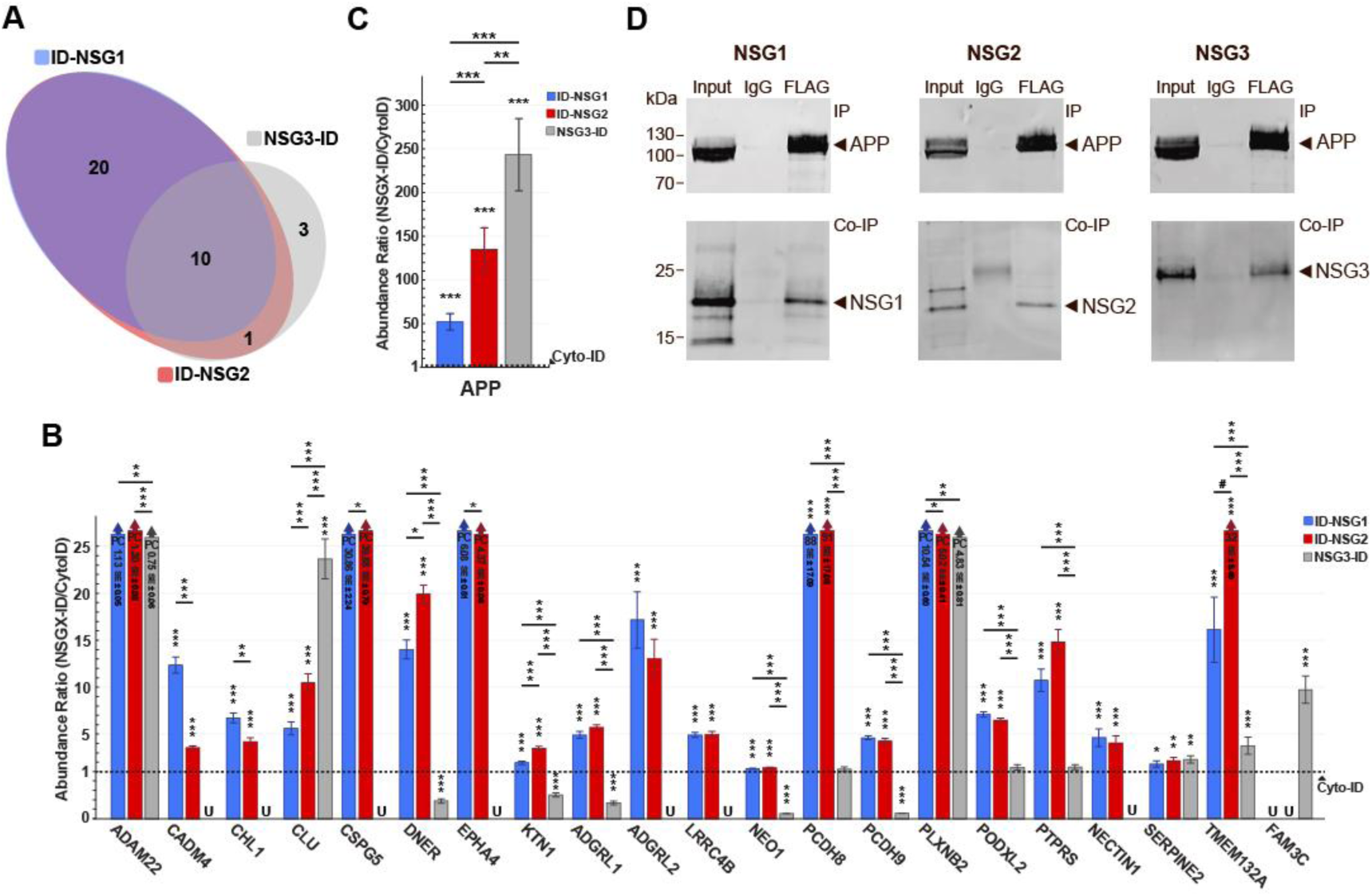
NSG1-3 interactome includes APP and multiple ADAM10 substrates. (A) Euler diagram showing the enrichment preference to NSGX-ID groups of ADAM10 substrates that are significantly represented in the NSGX-ID proteome list. Abundance ratios of NSGX-ID groups compared to Cyto-ID control (dashed line at “1”) show: (B) Subset of ADAM10 substrates enriched in across NSGX-IDs. (C) APP abundance across NSGX-ID group, comparison across experimental groups shows NSG3-ID biotinylated APP the most. (D) Co-IP of Flag tagged APP and probing for NSG family members shows interaction of APP with all NSG family members in transiently transfected HEK293 cells. For all bar graphs, “U” denotes undetected proteins in the corresponding NSGX-ID group. “PC” labeled bars show ratio and standard error (SE) in millions. Non-PC bars that have a large abundance ratio were restricted, ratio and SE stated inside bar. *# p < 0.1, *p < 0.05, **p < 0.01, ***p < 0.001*.

Due to the involvement of ADAM10 with Alzheimer’s Disease (AD)-related proteins, as well as a previous study showing NSG1 interaction with amyloid precursor protein (APP; Norstrom et al., 2010), we examined whether all NSG proteins could potentially interact with APP and APP-binding proteins. Figure 8C illustrates that, not only do NSG2 and NSG3 biotinylate APP to a greater extent than Cyto-ID, but they do so to a significantly greater extent than known binding partner NSG1 (Norstrom et al., 2010). Further, ID-NSG1/2 both biotinylated APP family members APLP1 and APLP2 in similar fashion to that of APP (Supplemental Figure 7), as well as a significant number of APP-binding proteins including members of the BRI2 and BRI3 family (ITSM2B and 2C, respectively; Supplemental Figure 7), which are well-studied proteins involved in amyloid-beta metabolism. Finally, we expressed FLAG-tagged APP-695 along with individual NSG1-3 family members in HEK-293 cells and used Co-IP to probe for physical interactions. IP using the Anti-FLAG antibody was able to successfully pull-down full-length APP in all conditions (Figure 8C, upper blots, right lanes) as well as to Co-IP each of the individual NSG1-3 proteins (Figure 8D, lower blots, right lanes). In contrast, use of an anti-IgG control antibody was not able to specifically IP APP nor any NSG family member. Together, these data augment the number of potential ADAM10 targets that may be differentially regulated by NSG family members (Overby et al., 2023), and specifically implicate all NSG family members in APP regulation.

## DISCUSSION

Characterizing the NSG interactome has the potential to clarify the obscure roles these proteins have in shaping neuronal function. Here, we found that NSG1 and NSG2 share a highly overlapping interactome, enriched for proteins involved in synaptic transmission, endosomal trafficking, and proteolytic processing—supporting a coordinated role in regulating receptor localization and neuronal signaling. In contrast, NSG3 displayed a partially overlapping but functionally distinct interactome, marked by preferential association with components of the endoplasmic reticulum and translational machinery. Additionally, all three NSG proteins showed previously unrecognized associations with presynaptic and inhibitory synaptic proteins, suggesting broader involvement in shaping both excitatory and inhibitory circuitry. Finally, our data provide support for an expanded role NSG family members in regulating proteolysis via associations with ADAM10, especially proteins involved in Alzheimer’s disease such as APP.

While NSG1-3 have all been implicated in regulation of synaptic function via AMPAR trafficking, interactome data offer insights into potentially unique roles between family members. For instance, while we and others have established interactions between NSG1 and NSG2 with the GLUA1 and GLUA2 subunits of AMPARs, the role of NSG3 remained unknown. Interactome data suggest that NSG3 may specifically interact with GLUA2 (Figure 5A), the predominately RNA-edited subunit that controls multiple AMPAR properties such as calcium permeability (Hume et al. 1991). These data also suggest that NSG2 may uniquely interact with GLUA3 and GLUA4-containing AMPARs (Figure 5A). Given the tightly regulated temporal and spatial patterns of AMPAR subunit expression (Zhu et al., 2000; Luh et al., 2009; Schwenk et al., 2014), future studies should explore whether individual NSG1-3 interactions have unique roles in regulating subpopulations of AMPARs similar to other AMPAR-binding proteins (Reimers et al., 2012; Herring et al., 2013). These findings may have significant implications in unique forms of plasticity where different AMPAR subunits play outsized roles especially during development (Huupponen et al., 2016; Luchkina et al., 2017).

Further, interactome data suggest possible mechanistic insights for how NSGs regulate AMPAR trafficking. Impairment of retromer function via knockdown of a key component of the complex blocks long-term potentiation (LTP; Temkin et al., 2017) which mimics interference with NSG1 function (Steiner et al., 2002; Alberi et al., 2005). Retromer helps sort recently endocytosed cargo between recycling and degradation pathways (Burd and Cullen, 2014), a function also ascribed to NSG1 (Debaigt et al., 2004; Steiner et al., 2002 and 2005). The current data suggests a synergistic, rather than redundant role for these two proteins, whereby NSGs and retromer work together to promote AMPAR recycling at synapses undergoing plasticity (Figure 7E). In addition, these data indicate that NSG1 and NSG2 are primarily involved in the “slow recycling” pathway, which is driven primarily by complexes CHEVI and FERARI alongside Rab11 and STX12 (reviewed in Jovic et al., 2010 & van der Beek et al., 2019). Complexes involved with the “fast recycling” of proteins from EE and SE to the PM include the EARP complex, VPS3 and RAB4, which were not identified in our study (reviewed in Jovic et al., 2010 & van der Beek et al., 2019). Interestingly, early studies of NSG1 identified Rab4 as a co-localizing protein, but recent studies failed to replicate these data, lending further evidence that NSGs are likely not involved in fast recycling pathways (Steiner et al., 2002; Yap et al., 2017). Moreover, MTCs have been characterized across different cell types and organisms, while NSGs are primarily expressed in neurons. This would indicate that NSGs are likely to act as auxiliary proteins for cargo recognition in the context of neuron specific protein trafficking, rather than playing a role in tethering vesicles.

Interactome data also potentially expands the role of NSGs in regulating synaptic function. First, while NSGs have not been implicated in regulating NMDARs, a previous report demonstrated the formation of a multi-protein complex between NSG3 (Calcyon) and Dopamine D1 receptors linked by PSD-95 (Ha et al., 2012). NMDARs are delivered and anchored within the post-synaptic density (PSD) by members of the DLG family, including PSD-95 and SAP102 (Niethammer et al., 1996; Washbourne et al., 2004; Lau and Zukin, 2007), and Electron Micrography (EM) studies have shown NSG1 localized at a subset of PSDs (Utvik et al., 2009). While ID-NSG1 and ID-NSG2 biotinylated all DLG family members to a greater degree than Cyto-ID, NSG3-ID biotinylation levels were significantly less than Cyto-ID (Figure 5C). It is intriguing that the obligate GRIN1 (GluN1) subunit of NMDARs (Monyer et al., 1992) was not detected in any group, suggesting steric hinderance of access of the TurboID protein for GluN1 subunits, technical limitations for detecting GluN1 peptides, or finally that NSGs may regulate unassembled NMDAR subunits prior to tetramerization to GluN1. Second, interactome data suggest possible roles within the presynaptic bouton, which enrichment was primarily driven by NSG1 and NSG2 interactomes. Indeed, we noted a great extent of colocalization of NSG1 and NSG2 at presynaptic sites. Others have reported and characterized specialized Golgi vesicles that deliver structural proteins to presynaptic sites (Maas et al., 2012), given the known enrichment of NSGs at the Golgi, also highlighted in these results, further studies will need to confirm whether NSGs may be involved in the sorting or trafficking of these Golgi vesicles. Additionally, NSG1 and NSG3 have been shown to be involved in the transcytosis of certain receptors from dendrites to axonal compartments. Data previously published from our lab has demonstrated that disrupting NSG2 expression reduces frequency of spontaneous EPSC, a measurement of presynaptic function (Chander et al., 2019). Further investigations will be needed to determine how NSGs could potentially be involved in formation, maintenance, or active neurotransmitter release at presynaptic sites.

While NSG3 has been reported in EM studies to be present in both postsynaptic and presynaptic sites PSD (Xiao et al., 2006; Negyessy et al., 2008), the current findings offer little support for a direct role of NSG3 in postsynaptic function. For example, the NSG3 interactome did now show overrepresentation of postsynaptic related pathways and few PSD proteins were significantly enriched in NSG3-ID groups. We also looked at potential biotinylation of spine apparatus (SA) proteins given NSG3 ER enrichment but failed to detect significant biotinylation of SYNPO, a known SA protein. In line with our findings, a recent proteomic study looking at subdomains of postsynaptic fractions did not identify NSG3 in their study, while both NSG1 and NSG2 were present in synaptic membrane fractions but not PSD fractions (Pandya et al., 2017). While NSG3 has not been shown to bind to AMPAR subunits, significant biotinylation of GLUA2 and TARPγ2/5 by NSG3-ID suggests possible role in AMPAR binding or potentially in trafficking TARPs independent of AMPARs.

NSG interactome and co-localization data also indicate a potential role in the trafficking of GABA_A_ receptors. We have previously shown that both NSG1 KO and NSG2 KO animals show perturbed diurnal activity, while NSG1 KO animals display enhanced anxiety and NSG2 KO animals show enhanced associative learning in a fear conditioning task (Austin et al. 2022, Zimmerman et al., 2024). Interestingly, recent studies have shown dynamic trafficking of GABA_A_ receptors in a sleep-wake dependent manner in the hippocampus (Wu et al., 2022) as well as long term changes of GABA_A_ receptors at inhibitory synapses in the amygdala as a result of associative fear learning (Kasugai et al., 2019). Further, many genetic perturbations that result in cognitive enhancement have a net effect of decreasing inhibition (Miyake et al., 2009; Moore et al., 2010; Murphy et al., 2004; Zhu et al., 2011). Thus, at least for NSG2 KO animals that display enhanced learning and memory (Zimmerman et al., 2024), one potential mechanism may be via interactions with GABA receptors, that ultimately leads to reduced inhibition with the loss of NSG2. GSEA also identified several pathways related to the ER, Ribosomes, and RNA binding of which NSG3-ID seemed to be the stronger driver of enrichment. As previously stated, ER related pathway enrichment was likely due to PTMs that NSG3 undergoes (Nilsson et al., 2009; Halim et al., 2013), which could be indicative of diverging trafficking roles of NSG3 compared to NSG1 and NSG2. Alongside ER, we also noted other gene expression related pathways that could indicate a novel function for NSG3, indeed NSG3 has been reported to indirectly affect protein expression upstream the Erk1/2 pathway (Xia et al., 2021) though not by interactions with RNA binding proteins. Similarly, in a more relevant model of pediatric pilocytic astrocytoma (PA), loss of NSG3 was associated with differential gene expression in PA tumors (Potter et al., 2008).

Potential flaws contained in the current study include previously demonstrated interactions that were not captured by proteomic analysis. For instance, GRIP1, SPARCL (Hevin), Clathrin Light Chain (CLTB), and neurotensin receptors (NTSR1-3, including SORT1) have all been demonstrated to associate with various NSG family members (Steiner et al., 2002 & 2005; Xiao er al., 2006; Kim et al., 2021; Overby er al., 2023). A plausible explanation for the absence of GRIP1 and CLTB, both of which were biotinylated by NSG family members, is that these proteins are relatively accessible to Cyto-ID mediated biotinylation and may therefore have been excluded during subsequent filtering steps. Indeed, GRIP1 is found as membrane bound and cytosolic pools (Hanley and Henley, 2010; Tan et al., 2015), The larger volumetric space occupied by Cyto-ID within cytosolic compartments, compared with membrane-restricted NSGX-ID constructs, could bias Cyto-ID toward labeling proteins that reside in both membrane-associated and cytosolic compartments. Additionally, although the human i3N model has been validated to generate cortical-like neurons (Wang et al., 2017), comprehensive proteomic profiling of this system has not yet been reported. This gap may contribute to the apparent absence of certain expected proteomic targets. Finally, because most NSG family studies have been conducted in rodent systems, species-specific differences that we did not directly assess may also underlie some of the divergences observed here. Supporting this possibility, we identified the human-specific adhesion molecule NLGN4X as enriched within NSG family proteomes.

## MATERIALS AND METHODS

### Cell Culture

All neuronal cultures were grown at the desired density directly on six-well plates (Corning) or in 24-well plates (Corning) on 12-mm coverslips (Electron Microscopy Sciences) coated with 0.1mg/mL poly-Ornithine (Sigma), 5µg/mL laminin (Corning) and 0.5µg/mL Matrigel (Corning). HEK293T cells were grown in standard 10cm tissue culture plates (Sarstedt). Neurons were transfected using Lipofectamine LTX with Plus Reagent (Thermo Fisher Scientific) as per manufacturer’s recommendation.

HEK293T cells were cultured as a monolayer at 37°C under 5% CO_2_ in Dulbecco’s Modified Eagle Medium (DMEM; Corning) media supplied with 10% fetal bovine serum (FBS; Gibco) and 1% Penicillin-Streptomycin (Pen Strep; Gibco). HEK293T cells grown to 90% confluency and were passaged using 0.05% Trypsin-EDTA (Gibco).

Primary hippocampal neurons were cultured from neonatal (P0-P1) C57BI/6J pups essentially as described in (Chander et al., 2019). In short, brains were removed, and the hippocampus was dissected in ice-cold Hank’s Balanced Salt Solution (HBSS; Sigma) supplemented with 20% FBS and 4.2mM NaHCO_3_ (Sigma), 1mM HEPES (Sigma) at pH 7.4. Hippocampi were digested in 0.25% Trypsin-EDTA (Gibco) for 10 minutes at room temperature (RT) and dissociated using fire polished Pasteur pipettes of decreasing diameter in ice-cold HBSS containing 1500U DNase (Sigma). Neurons were pelleted, resuspended in plating media, and plated at a density of 4-5 x 10^5^ cells per coverslip. Cells were left to adhere for 15 minutes, followed by addition of 0.5mL of plating media composed of Neurobasal (Gibco) supplemented with 1x B27 (Gibco), 1x GlutaMAX (Gibco), 1% Pen Strep, and 5% FBS for 24 hours. FBS was eliminated from this media after 24 hours and again replaced after 48 hours supplemented with 4µM cytosine 1-β-D arabinofuranoside (Ara-C; Sigma). Hereon, half the volume of media was replaced every week with fresh media without Ara-C and FBS. All animal procedures were performed in accordance with an approved protocol by the University of New Mexico-Health-Science Institutional Animal Care and Use Committee.

i^3^N iPSCs were cultured as a monolayer at 37°C under 5% CO_2_ in mTeSR Plus medium supplied with 1x mTeSR Plus Supplement and passaged at 90% confluency using 0.5mM EDTA (Sigma). i^3^N iPSCs were differentiated to i^3^Ns using a previously described protocol (Wang et al., 2017, Fernandopulle et al., 2018). In short, i^3^N iPSCs were incubated with 20nM doxycycline (Sigma) for 3 days at a density of 2.0–2.5 × 10^6^ cells/well in six-well plates in supplemented mTeSR Plus medium and 10μM Y-27632 (Abcam). The full volume of medium was changed daily. These pre-differentiated i^3^N precursor cells were dissociated, counted, and plated at a density of 5-9 x 10^4^ cells per cm^2^ in media mixture composed of 50% DMEM/Nutrient Mixture F-12 Ham (Sigma) and 50% Neurobasal supplemented with 10ng/mL BDNF (Prepotech), 10ng/mL GDNF (Prepotech), 10ng/mL NT3 (Prepotech), 1μM cAMP (Sigma), 200μM ascorbic acid (Sigma), 0.5µg/mL laminin, and x1 B27. i^3^Ns were co-cultured alongside mouse astrocytes obtained from P0-P1 using similar methods described above and Ara-C was added at day 3 from co-culture set up. Half of the medium was replaced on day 7 and again on day 14, and the medium volume was doubled on day 21. Thereafter, one-third of the medium was replaced bi-weekly until the cells were used for experiments.

### DNA Plasmids

A total of four different TurboID plasmids were cloned using standard restriction enzymes, PCR amplification, and NEBuilder HiFi DNA Assembly kit (NEB) using manufacturer recommendations. TurboID constructs were cloned into a pFCK plasmid backbone downstream of the Synapsin-1 promoter, where restriction enzymes BamHI and EcoRI were used to linearize the backbone plasmid. We generated primers for PCR amplification containing overlap regions to the pFCK backbone that also targeted the open reading frame (ORF) of V5-TurboID-NES_pCDNA3 (Addgene #107169) to generate Cytoplasmic-TurboID (Cyto-ID). Similarly, we created primers that would target the ORF of human NSG1 (AK312288.1), human NSG2 (AK312359.1), or human NSG3 (AF225903.1) while promoting correct N-terminal or C-terminal plasmid incorporation. The linker peptide sequence GGSGGSGGSGG was designed into the constructs between each NSG1-3 and V5-TurboID. Constructs were verified by sanger sequencing confirmation of ORF V5-TurboID-NSG1, V5-TurboID-NSG2, and NSG3-V5-TurboID. NSG3-mNGreen plasmid was used for ICC experiments where endogenous NSG3 was undetected by antibodies. pcDNA-FLAG-NSG3 (Calcyon) was a generous gift from Clare Bergson.

### Lentivirus Production

Preparations of lentivirus were done in-house using a second-generation packaging system according to publish methods (Binder et al., 2020). Briefly, HEK293T cells were grown as described above and transfected at 80% confluency using a Calcium Phosphate protocol (Kingston et al., 2003) using 10μg of assembled TurboID plasmid, 5μg of the second-generation packaging plasmid psPAX2 (Addgene #12260) and 2.5μg of viral envelope protein VSV-G plasmid pMD2.G (Addgene #12259). 3-4 days post-transfection, medium was collected, centrifuged for 5 minutes at 300 x g and filtered through a 0.45-μm filter (Millipore Sigma). Lentiviral particles were concentrated via ultracentrifugation of medium at 25,000 x g for 2 hours at 4°C on a bed of 20% sucrose (Sigma) in phosphate-buffered saline (PBS; VWR). Pelleted virus was resuspended in 250µL-300µL of ice-cold Neurobasal, left overnight in a rotary agitator at 4°C, and aliquoted for storage at −80°C until use.

### Immunocytochemistry

Immunocytochemistry (ICC) protocol used has been modified from previously described Larsen et al., 2016. In short, coverslips were fixed in 3.2% paraformaldehyde (PFA; Electron Microscopy Sciences) for 10 minutes, washed x3 with PBS, permeabilized with 0.2% Triton X-100 (Amresco) for 5 minutes, washed x3 with PBS, and blocked in 10% donkey serum (DS; Sigma) for 30 minutes. Primary antibody diluted in 5% DS and coverslips were incubated for 4 hours at RT or overnight at 4°C. Primary antibodies used: mouse anti-V5 (1:3000; Bio-Rad), goat anti-NSG1 (1:2000; Invitrogen), rabbit anti-NSG2 (1:500; Abcam), rabbit anti-NSG3 (1:1000; Invitrogen), chicken anti-MAP2 (1:2000; Novus Biologicals), rabbit anti-VPS26B (1:500; Novus Biologicals), guinea pig anti-SYN1 (1:1000; Sysy), rabbit anti-GluN2B (1:500; Phosphosolutions), rabbit anti-tomm20 (1:500; Novus), and rabbit anti-calreticulin (1:500; Cell Signaling), anti-GABRB2 (1:500; Kind gift from Dr. Clare Bergson). After primary antibody incubation, cells were washed x3 with PBS and then incubated for 1 hour at room temperature with secondary antibodies in 5% DS. Conjugated secondary antibodies used: goat anti-chicken CF405M (1:500; Millipore Sigma), donkey anti-guinea pig DL405 (1:500; Jackson ImmunoResearch), streptavidin AF488 (1:5000; Jackson ImmunoResearch), donkey anti-chicken AF488 (1:2000; Jackson ImmunoResearch), donkey anti-goat AF488 (1:2000; Invitrogen), donkey anti-rabbit AF488 (1:2000; Invitrogen), donkey anti-mouse DL550 (1:1000; Invitrogen), donkey anti-rabbit DL550 (1:1000; Invitrogen), donkey anti-mouse AF647 (1:1000; Jackson ImmunoResearch), donkey anti-goat AF647 (1:1000; Jackson ImmunoResearch), and donkey anti-rabbit AF647 (1:1000; Jackson ImmunoResearch). Finally, coverslips were washed x3 with PBS and mounted on glass slides using Fluoromount-G (Southern Biotech).

### Confocal Imaging

Confocal z-stack images were acquired on the Zeiss LSM800 airyscan confocal microscope using the 63x/1.40NA oil-immersion objective. Sequential frame acquisition was set to acquire an average of 14 planes per stack at 16 bit and a resolution of 1216 x 1216. Channel gain settings were optimally adjusted to minimize saturation of punctae and were maintained across experimental groups. Unmodified images were used for all analyses and linear scaling was applied on images only for presentation purposes using ZEN 3.8.

### Western Blots & Immunoprecipitations

Cells were quickly washed x3 with ice-cold phosphate-buffered saline (PBS; VWR). Following washes, 500µL of RIPA lysis buffer composed of 50 mM Tris buffer (Bio-Rad) at pH 8.0, 150 mM NaCl (RPI), 0.1% SDS (Amresco), 0.5% sodium deoxycholate (Sigma), 1% Triton X-100, and x1 protease inhibitor tablet (Pierce), was added to each well of a 6-well plate, cells were then scraped and collected. Lysed cells were rotated at 4°C for 20 minutes and debris was pelleted by centrifugation at 13,000 x g for 15 minutes. Supernatant was collected and stored at −80°C until use. Protein lysate concentration was quantified using the BCA Protein Assay Kit (Pierce). Protein samples were denatured in x1 NuPAGE sample reducing buffer (Invitrogen) and x1 NuPAGE LDS sample buffer (Invitrogen) and boiled for 20 minutes at 95° C. In a Criterion blotter system (Bio-Rad), 10-25µg of protein sample was loaded per well and separated on an 10% SDS-PAGE gel and transferred to a PVDF membrane (Li-COR) and stained using standard immunoblotting protocols (Mahmood and Yang, 2012). Primary antibodies used: mouse anti-V5 (1:2000), mouse anti-NSG1 (1:1000; Santa Cruz), rabbit anti-NSG2 (1:500; Abcam), mouse anti-NSG3 (1:1000; Santa Cruz), and rabbit anti-APP (1:2000; Abcam). Conjugated secondary antibodies used: goat anti-mouse horseradish peroxidase (HRP) (1:10,000; Jackson ImmunoResearch), goat anti-rabbit HRP (1:10,000; Jackson ImmunoResearch), and streptavidin HRP (1:20,000; Jackson ImmunoResearch). Blots stained for biotinylated proteins using streptavidin-HRP staining were blocked and incubated with 3% bovine serum albumin (BSA; Sigma) in PBS 0.1% Tween-20 (PBST; Amresco), all other staining was incubated in and blocked with 5% milk (Bio-Rad) in PBST. HRP signal was developed with SuperSignal West Pico PLUS chemiluminescent substrate (Thermo Fisher Scientific) as per manufactures recommendation. All blots were scanned using the ChemiDoc MP Imaging System (Bio-Rad).

For co-immunoprecipitation, whole mouse brains were dissected and frozen on dry-ice before lysing with 1mL/ Half brain of Co-IP lysis buffer composed of 25mM Tris at pH 7.4, 150mM NaCl, 1mM EDTA, 5% Glycerol (JTBaker), 1% NP-40 (Sigma), x1 protease inhibitor tablet, and x1 phosphatase inhibitor (Sigma). Sample lysate was incubated with 10µg of target antibody overnight in a rotary agitator at 4°C. 25µL of A/G Magnetic Beads (Pierce) were washed 2x with Wash buffer composed of 50mM Tris at pH 7.5, 150mM NaCl, and 0.05% Tween-20, and pelleted with magnetic rack at all steps. A/G beads were then incubated with the antibody-sample lysate mixture for 1 hour in a rotary agitator at RT. Next, supernatant was separated and saved for quality control. Collected A/G beads were washed 3x with 500µL wash buffer and 1x with 500µL Ultrapure H_2_O. To elute proteins, 100µL of 0.1M Glycine (RPI) at pH 2.0 was added to A/G beads for 10 minutes in a rotary agitator at RT, supernatant was saved and neutralized with 15µL of 1M Tris at pH 8.0. Protein eluate was prepared for immunoblotting experiments as described above.

### Proximity Dependent Labeling with TurboID and Enrichment of Biotinylated Proteins

Proximity dependent protein labeling and enrichment is based on a modified published protocol (Cho et al., 2020), optimized and scaled for the i3N model system. i^3^Ns and mouse hippocampal neurons were transduced with lentiviral particles 7 days prior to experiments. 50µM of D-biotin (GoldBio) was supplied to the media approximately 3 hours before sample collection. Biotin labeling was stopped by placing the cells on ice and washing x3 with ice-cold PBS and subsequent lysing for immunoblotting and proteomic experiments or fixing for ICC experiments.

For biotinylated protein enrichment, we used 600ug of RIPA protein lysate and 0.6mg of streptavidin magnetic beads (Pierce), pelleted beads with magnetic rack, and performed all steps at RT unless otherwise stated. Magnetic beads were washed x2 with 1mL RIPA lysis buffer and incubated with protein lysate overnight in a rotary agitator at 4°C. After incubation, supernatant was saved for quality control and beads were washed x2 with 1mL of RIPA lysis buffer for 2 minutes, x1 with 1mL of 1M KCl (Sigma) for 2 minutes, x1 with 1mL of Na_2_CO_3_ (Sigma) for 10 seconds, x1 with 1mL of 2M Urea (Sigma) in 10 mM Tris-HCl at pH 8.0 for 10 seconds, and x2 with 1mL of RIPA lysis buffer for 2 minutes. After all washing steps, magnetic beads were resuspended in 500µL of RIPA lysis buffer, 5% of resuspended beads were saved for quality control, and remainder was transferred on-ice to core facility for protein digestion.

### Protein Digestion

Proteins bound to magnetic beads were washed x1 with 50μL of 50mM Tris-HCl at pH 7.5 and x2 with 50μL of 2M Urea in 50mM Tris buffer at pH 7.5. Magnetic beads were then incubated with 26 μL of 2M Urea in 50 mM Tris-HCl at pH 7.5, containing 1mM DTT and 90ng Trypsin at 25°C for 1 hour while shaking at 1,000 RPM. The supernatant was then removed and transferred to a new tube. Next, magnetic beads were washed x2 with 15μL of 2M Urea in 50mM Tris buffer at pH 7.5 and the wash solutions were combined with the supernatant from the previous step. Peptide supernatant was supplemented with additional DTT for a final concentration of 4mM to reduce disulfide bonds and incubated for 30 minutes with shaking at 1,000 RPM. Then, supplemented with iodoacetamide to a final concentration of 10 mM for alkylation and incubated for 30 minutes while shaking at 1,000 RPM in the dark. Finally, an additional 100ng of trypsin was added to the sample and digestion was ran overnight at 37°C with shaking at 200 RPM. After the overnight digestion, the solution was acidified by trifluoroacetic acid to approximately pH 3.0 and desalted by C18 ZipTip (ZTC18S096, Millipore Sigma) by following procedures in the product manual. The eluent from the ZipTip was dried in a SpeedVac vacuum concentrator and reconstituted with 6μL of 0.1% formic acid, of which 1.5μL was used for LC-MS/MS experiment. All protein samples were processed in parallel.

### Proteomics by LC-MS/MS

LC-MS/MS analysis was performed on a Q Exactive Classic mass spectrometer (Thermo Fisher Scientific, San Jose, CA) equipped with an EasySpray ion source. Peptides were eluted from an Acclaim Pepmap 100 trap column (75 micron ID x 2 cm; Thermo Fisher Scientific) onto an Acclaim PepMap RSLC analytical column (75 micron ID × 25 cm; Thermo Fisher Scientific) using mobile phases composed of 0.1% formic acid in water (A) and 0.1% formic acid in acetonitrile (B), and a gradient of 2-40% B over 127 minutes, 40-90% B over 3 minutes, holding 90% B for 7 minutes, 90-2% B over 2 minutes, and lastly equilibrating at 2% B for 13 minutes. Flow rate was 300nL/min using a Dionex Ultimate 3000 RSLCnano System (Thermo Fisher Scientific). Data dependent scanning was performed using a survey scan at 70,000 resolution, scanning mass/charge (m/z) 375-2000, automatic gain control (AGC) target of 3e6 and a maximum injection time (IT) of 10 ms, followed by higher-energy collisional dissociation (HCD) tandem mass spectrometry (MS/MS) at 30 normalized collision energy (nce), of the 15 most intense ions at a resolution of 17,500, an isolation width of 1.7 m/z, an AGC of 2e5 and a maximum IT of 75 ms. Dynamic exclusion was set to place any selected m/z on an exclusion list for 30 seconds after a single MS/MS. Isotopes were excluded from MS/MS. Technical duplicates were performed for each sample.

### Proteomics Data Analysis

Tandem mass spectra were searched against UniProt Homo Sapiens database (revi_2023_10_26) using Thermo Proteome Discoverer v2.5 (Thermo Fisher Scientific, San Jose, CA) considering fully tryptic peptides with up to 2 missed cleavage sites and static modification of cysteine carbamidomethylation (57.021 Da). Variable modifications considered during the search included methionine oxidation (15.995 Da), and protein N-terminal acetylation (42.011 Da). Proteins were identified with precursor mass tolerance of 10 ppm, fragment mass tolerance of 0.02 Da, and 1% FDR. Intensity based precursor abundance was used for label-free quantitation. Features detected in a minimum of 50% of replicates in the same experiment group were considered for quantitation. Normalization was performed based on total peptide amount. For statistical analysis, individual protein based one-way ANOVA analysis with Benjamini-Hochberg correction on the p-value was performed using Thermo Proteome Discoverer v2.5. Proteins with a fold change equal to or greater than 2 and adjusted p-value less than 0.05 were considered differentially expressed between groups.

## SUPPLEMENTAL FIGURES

**Supplemental Figure 1:**
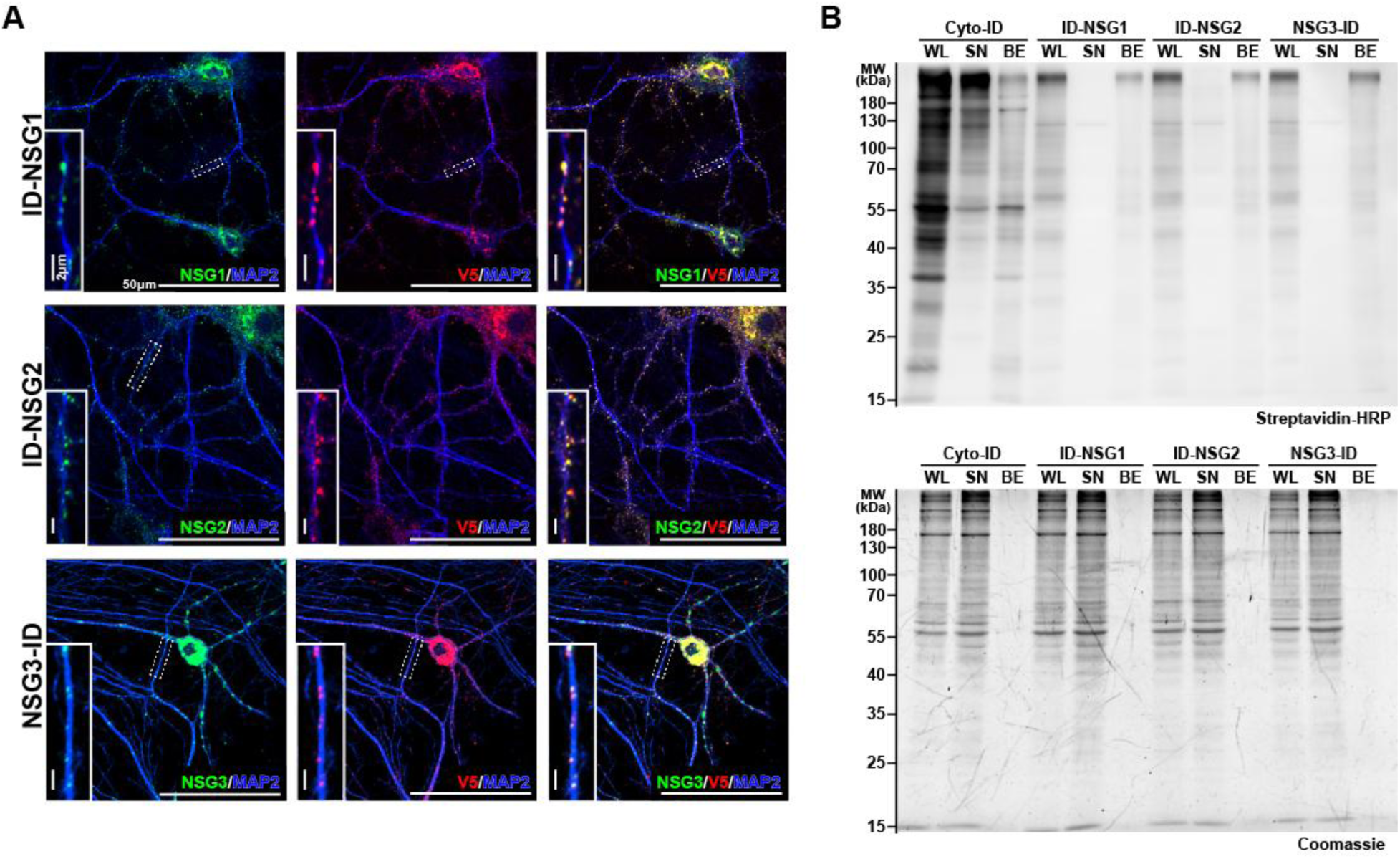
Correct expression of TurboID constructs and pulldown of biotinylated proteins. (A) ICC labeling of each TurboID-fusion protein demonstrates expected colocalization of signal using antibodies directed to endogenous NSG protein and V5 epitope tag. Scale bars, 50µm and 2µm. (B) Western blot analysis displaying levels of biotinylated proteins (top) from each step of the streptavidin-conjugated magnetic bead pulldown. For each group we loaded a known amount of protein for both whole neuronal lysate (WL) and supernatant after bead incubation (SN), an unknown amount of eluted proteins from a small portion of beads (BE) was also loaded per group. We detected biotinylated proteins in all WL and BE lanes and significant depletion in all SN lanes except for the Cyto-ID group. The second blot (bottom) shows total protein stain for each lane.

**Supplemental Figure 2:**
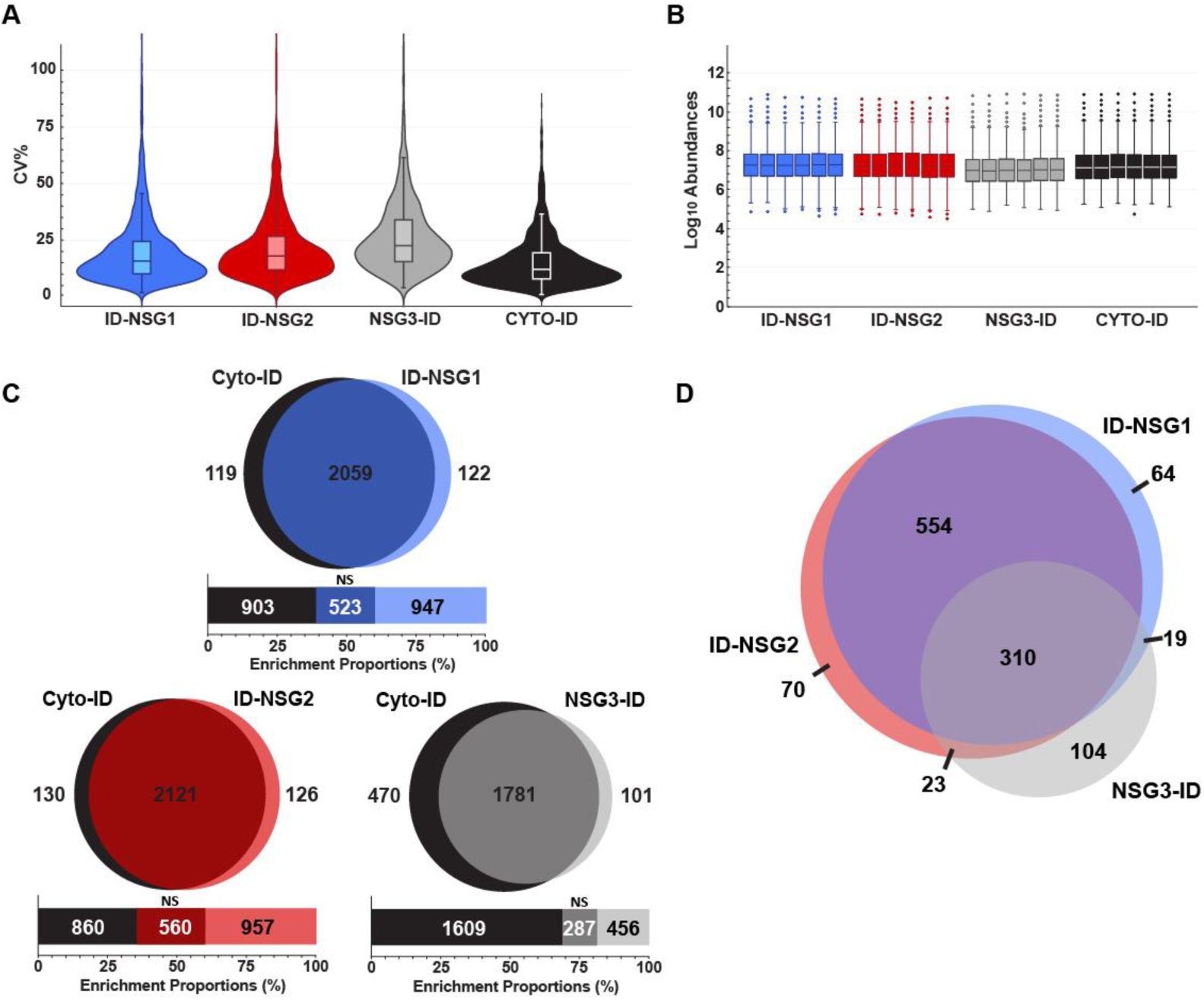
Proteomic quality and distribution across NSGX-ID constructs. (A) Violin distribution and boxplot graphs of coefficient of variation percentages (CV%) of all proteins within each NSGX-ID and Cyto-ID groups. Approximate medians: ID-NSG1 14.7%, ID-NSG2 17.8%, NSG3-ID 21.6%, and Cyto-ID 12.5%. (B) Boxplot of log_10_ protein intensities within each group across all technical replicates used for LC-MS. (C) Euler diagrams showing the overlap between the identified proteins within NSGX-ID groups and Cyto-ID. The compound bar graph indicates the number of proteins found to be significantly enriched (see methods) in both the NSGX-ID group (light blue, light red, or light gray) and Cyto-ID (black), the remaining group (dark blue, dark red, or dark gray) are the not significant (NS) proteins. (D) Euler diagram showing the overlap of the significantly biotinylated proteins of each NSGX-ID vs Cyto-ID comparison, where ID-NSG1 and ID-NSG2 have the greatest overlap and NSG3-ID has the greatest divergence.

**Supplemental Figure 3:**
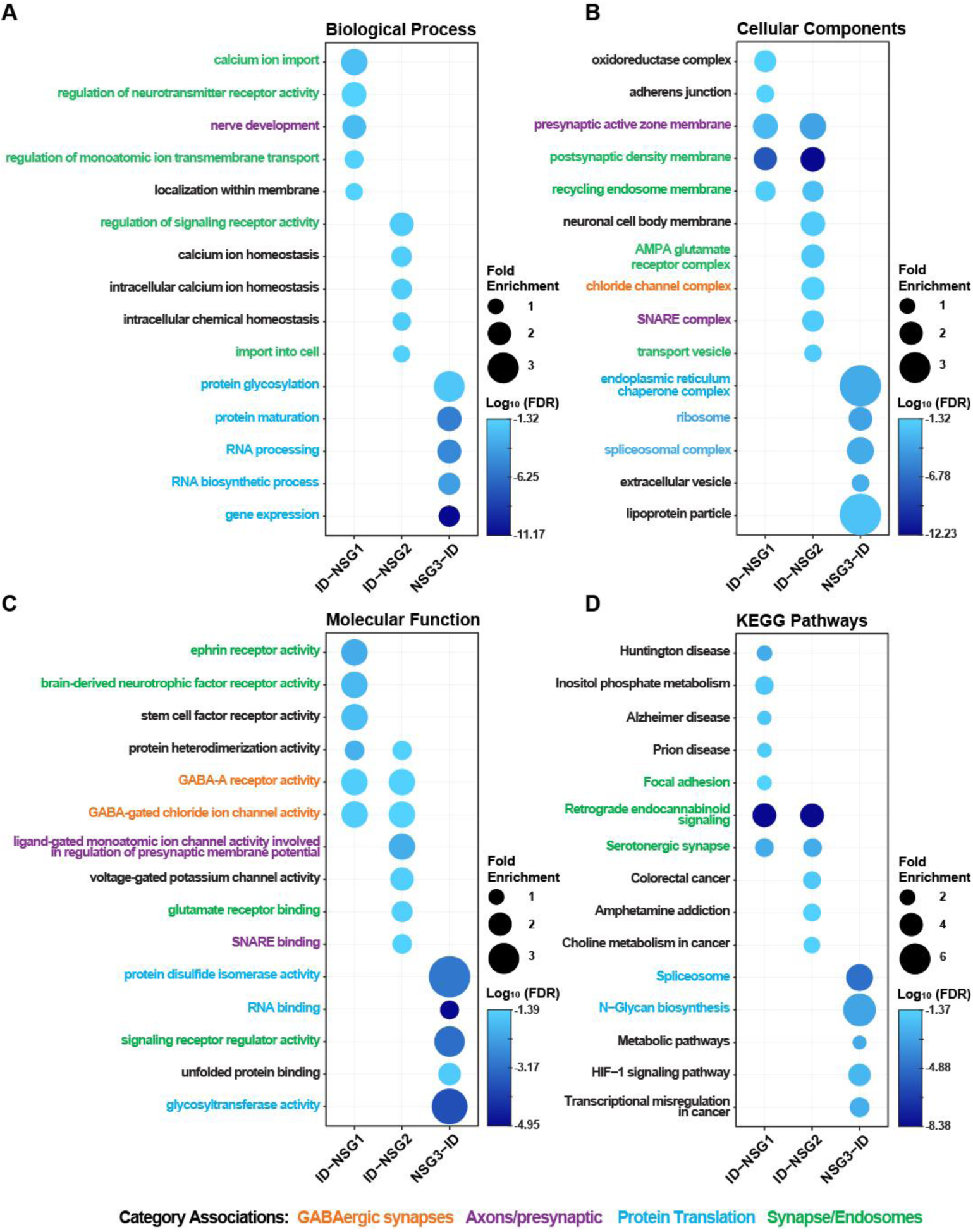
Distinct GO and KEGG Pathway enrichment profiles identify functional divergence among NSG1-3. (A-C) Gene Ontology and (D) KEGG Pathways GSEA of each NSGX-ID interactome (947, 957, and 456 proteins for NSG1-3, respectively) was done to identify diverging functional roles. Color coding of pathway text labels indicates known (green) versus novel (orange, purple, turquoise) functional associations, with turquoise denoting terms primarily enriched in the NSG3-ID dataset. NSG3-ID exhibited distinct enrichment for endoplasmic reticulum and protein translation-related pathways. ID-NSG1 and ID-NSG2 showed substantial overlap in their enrichment profiles, particularly among putative novel synaptic and endosomal categories. Bubble size corresponds to fold enrichment and pseudo-colored scale reflects the log₁₀(FDR).

**Supplemental Figure 4:**
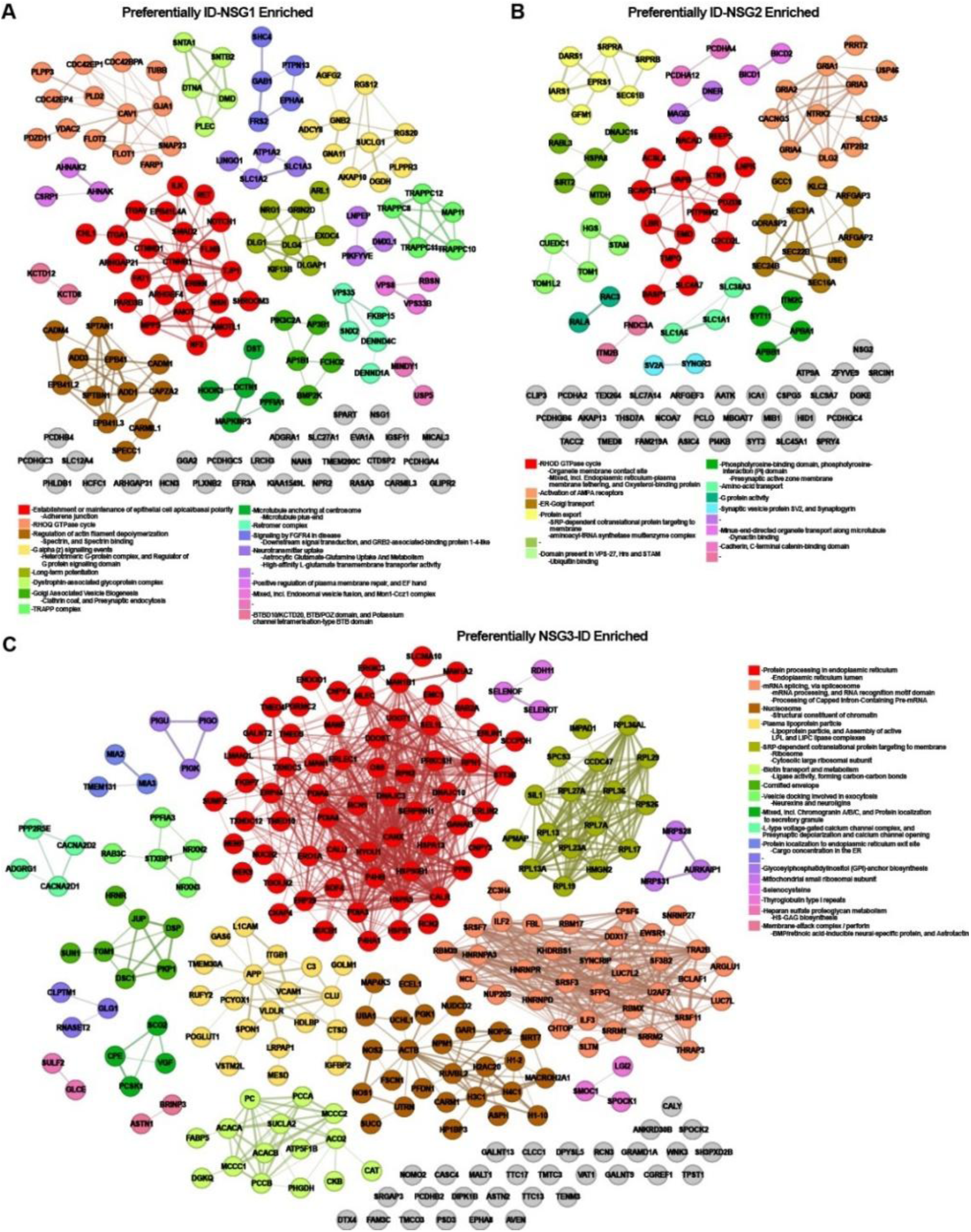
Protein-Protein interactions networks of NSG1-3 preferentially enriched proteins reveal potential NSG specific functions. (A-C) Protein-protein interaction (PPI) networks of specifically enriched proximity-labeled proteins for each NSGX-ID construct were generated using STRING database confidence scores. Proteins included in each network were significantly enriched preferentially in only one of the NSGX-ID groups. Each node represents a protein, and edge connections reflect STRING-derived confidence-based interactions. Node colors correspond to functional clusters identified using Markov Clustering (MCL). Annotated functional categories for each cluster are shown below each network. These interaction maps illustrate the modular architecture of each NSG construct’s interactome, with NSG3-ID revealing extensive clustering around ER and protein processing functions, while ID-NSG1 and ID-NSG2 exhibit more discrete clusters, including synaptic vesicle cycling, SNARE function, and disease-relevant pathways.

**Supplemental Figure 5:**
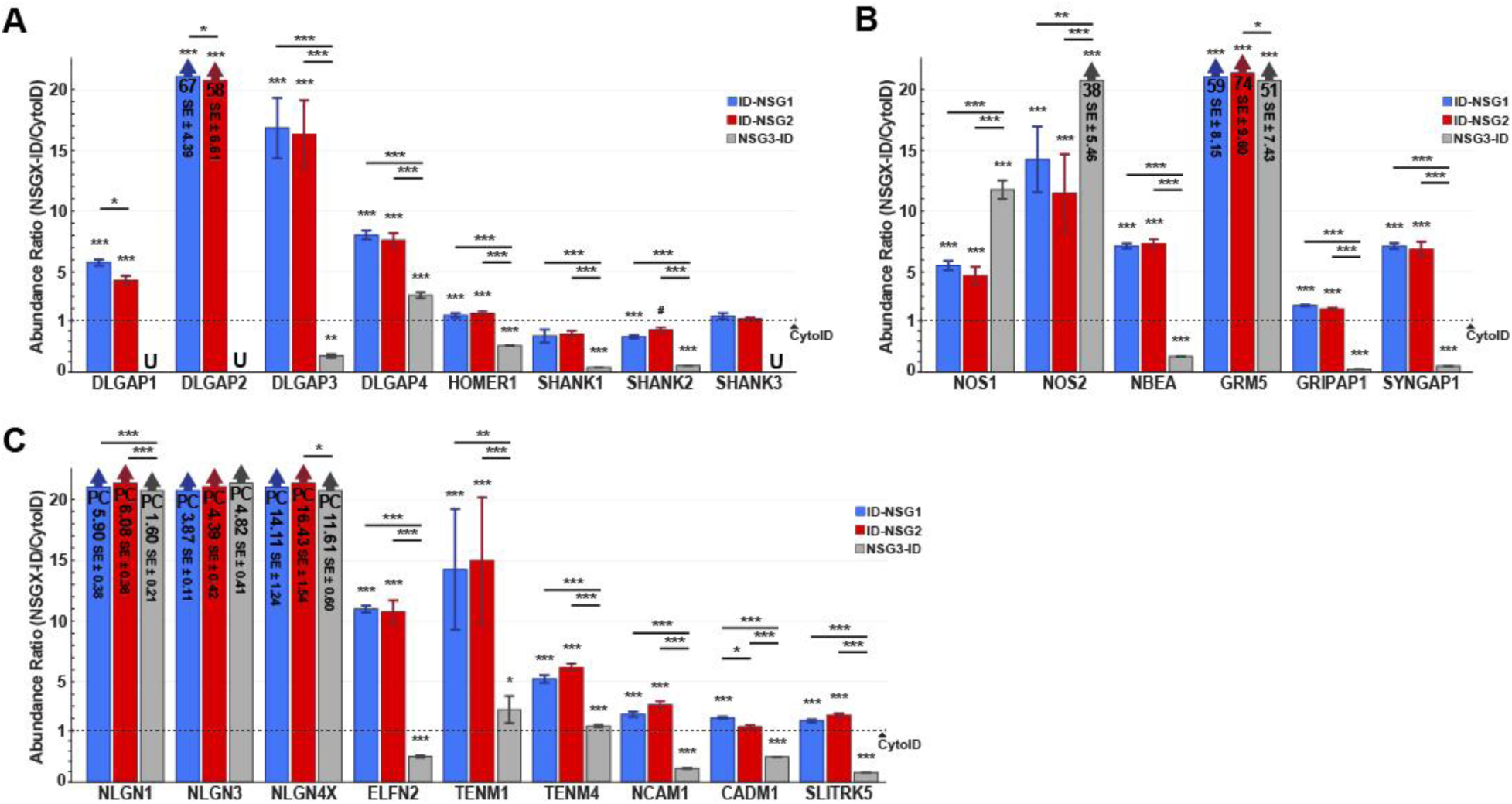
NSG1-3 potentially interact with proteins across a broad range of functional synaptic roles. Abundance ratios of NSGX-ID groups compared to Cyto-ID control (dashed line at “1”) show: (A) known postsynaptic scaffolding proteins, of which DLGAP1-4 are preferentially enriched in NSGX-IDs compared to other well-known scaffold proteins. (B) Various proteins involved in different forms of synaptic signaling. (C) Synaptic adhesion proteins that play a role in synaptic stability and signaling via secondary interactions. For all bar graphs, “U” denotes undetected proteins in the corresponding NSGX-ID group. “PC” labeled bars show ratio and standard error (SE) in millions. Non-PC bars that have a large abundance ratio were restricted, ratio and SE stated inside bar. *# p < 0.1, *p < 0.05, **p < 0.01, ***p < 0.001*.

**Supplemental Figure 6:**
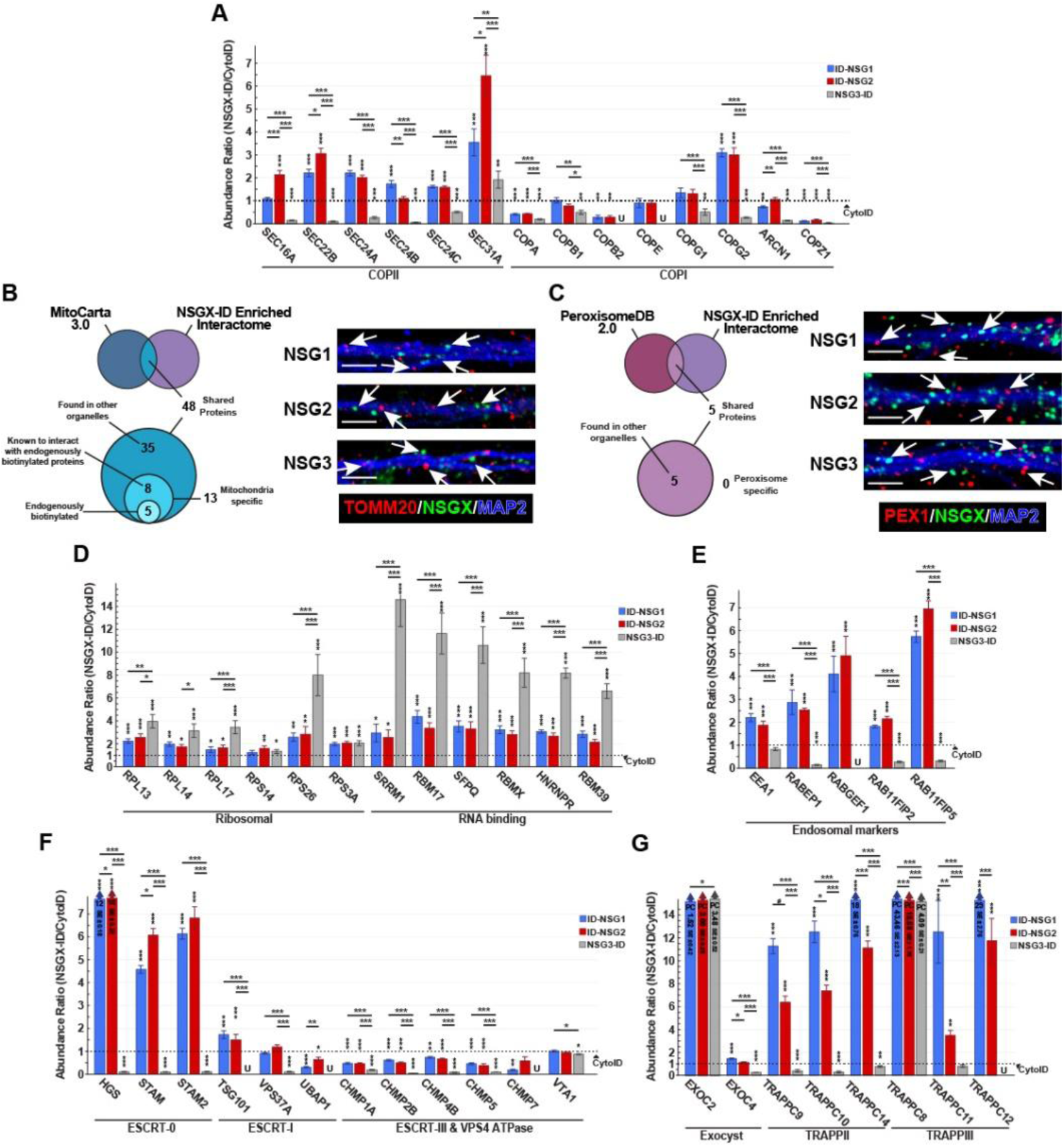
NSG1-3 interact with specific proteins involved in sorting and trafficking. Abundance ratios of NSGX-ID groups compared to Cyto-ID control (dashed line at “1”) show: (A) significant biotinylation by ID-NSG1 and ID-NSG2 of COPII coat subunits compared to COPI. Comparison of NSGX-ID enriched proteins with (B) MitoCarta 3.0 or (C) PeroxisomeDB 2.0 showed few organelle specific markers enriched within our NSG family interactome and complementary confocal images of (B) mitochondria marker TOMM20 or (C) peroxisome marker PEX1 lacking colocalization (full arrow) with any endogenous NSG1, NSG2, or transfected NSG3-mNGreen in cultured mouse neurons. Scale bars are 10 µm. (D) NSG3-ID preferentially enriched proteins involved in Ribosomal and RNA binding and processing. (E) Early endosome markers. (F) Enrichment of only ESCRT0 subunits by ID-NSG1 and ID-NSG2, no significant enrichment in other ESCRT complexes. (G) Exocyst and TRAPPII & III subunits enriched in NSGX-ID proteomes. For all bar graphs, “U” denotes undetected proteins in the corresponding NSGX-ID group. “PC” labeled bars show ratio and standard error (SE) in millions. Non-PC bars that have a large abundance ratio were restricted, ratio and SE stated inside bar. *# p < 0.1, *p < 0.05, **p < 0.01, ***p < 0.001*.

**Supplemental Figure 7:**
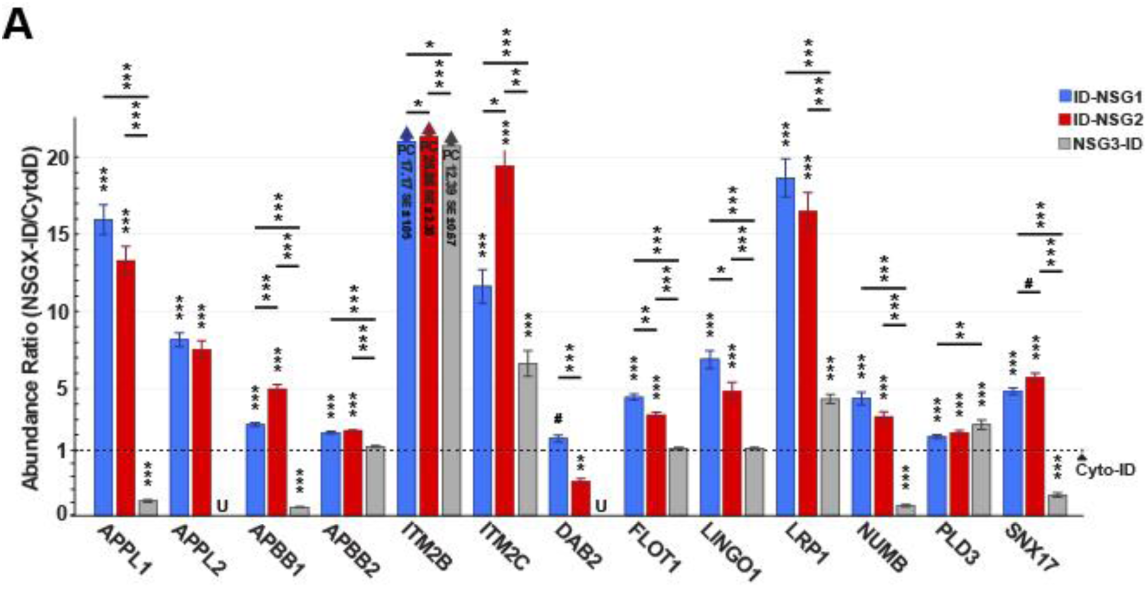
APP family members and known APP-binding proteins. Abundance ratios of NSGX-ID groups compared to Cyto-ID control (dashed line at “1”) show: (A) significant biotinylation by ID-NSG1 and ID-NSG2 of many APP family member proteins and APP-binding proteins. For all bar graphs, “U” denotes undetected proteins in the corresponding NSGX-ID group. “PC” labeled bars show ratio and standard error (SE) in millions. Non-PC bars that have a large abundance ratio were restricted, ratio and SE stated inside bar. *# p < 0.1, *p < 0.05, **p < 0.01, ***p < 0.001*.

